# cAMP responsiveness determines follicular ovulatory fate downstream of the LH receptor in the cloudy catshark

**DOI:** 10.64898/2026.04.10.717848

**Authors:** Ryotaro Inoue, Takanori Kinugasa, Keigo Nagasaka, Kotaro Tokunaga, Shigeho Ijiri, Susumu Hyodo

## Abstract

The number of offspring produced per reproductive cycle is generally explained by a luteinizing hormone receptor (LHR)-threshold model, in which only follicles expressing sufficient levels of LHR respond to the LH surge and proceed to ovulation. Using the cloudy catshark as a model, we reveal a distinct mechanism that determines the fate of ovulatory (F1) and non-ovulatory (F2) follicles independently of LHR abundance. The cloudy catshark produces only two eggs per reproductive cycle by ovulating a pair of F1 follicles from its single ovary. Both F1 and F2 follicles express comparable levels of *lhr*, indicating that both retain the capacity to receive the LH signal. Nevertheless, LH stimulation selectively activated ovulation-associated responses in F1 follicles. These included progesterone production via *star2* upregulation, as well as transcriptional programs including RUNX transcription factors, prostaglandin signaling (*ptgs2*, *ptger1*), and matrix metalloproteinases. Moreover, direct activation of cAMP pathway, bypassing LHR activation, induced progesterone production in F1 but not F2 follicles, indicating that the critical difference in ovulatory responsiveness lies downstream of cAMP. These results demonstrate that follicular ovulatory fate is determined, at least in part, by follicle-specific transcriptional responses to cAMP, rather than by LHR abundance alone, challenging the prevailing LHR-threshold model of ovulatory regulation in vertebrates.

## 1. Introduction

Reproductive strategies vary widely among animal lineages, ranging from K-strategists that typically produce a small number of high-investment offspring to r-strategists that release large numbers of low-investment eggs. For example, humans typically produce a single offspring per reproductive cycle requiring prolonged parental investment (Zeleznik, 2004), whereas many fish release thousands or even millions of eggs during a single spawning event (Barneche et al., 2018). Birds and reptiles possess hierarchical ovaries, in which follicles at different developmental stages are ovulated sequentially, yielding clutches ranging from a single egg to several tens of eggs per reproductive cycle (Etches and Petitte, 1990). Across these lineages, reproductive output is ultimately constrained by the number of follicles that develop and acquire ovulatory competence within each reproductive cycle.

In bony vertebrates, ovulation is regulated by the luteinizing hormone (LH) surge (Ando, 2016), and the expression of luteinizing hormone receptor (LHR) is considered a key determinant of whether a follicle can ovulate in response to the LH surge. This mechanism has been extensively studied in mammals and birds, in which ovulation number is typically low and tightly regulated. In mammals, at the onset of the reproductive cycle, a subset of primordial follicles initiate gonadotropin-dependent development (Longo et al., 2025; Zeleznik, 2004). With the progression of development, most follicles are arrested in their maturation, while the selected dominant follicle(s) acquire LHR expression in the follicular granulosa cells and proceed to ovulation in response to the LH surge (Matsuda et al., 2012). By contrast, in birds, follicles at vitellogenic stages progressively express LHR during development (Johnson et al., 1996). Both preovulatory and non-preovulatory follicles respond to the LH surge by secreting progesterone (P4) (Johnson et al., 2002a) and increasing protease activity associated with follicle rupture (Jackson et al., 1993). These responses are more pronounced in preovulatory F1 follicles, which exhibit higher levels of LHR expression than non-preovulatory follicles. This has led to the proposal that ovulation requires a threshold level of LHR expression (Liu et al., 2022a). Although the processes of follicular development differ between birds and mammals, ovulatory competence in both systems can be explained by a common framework in which LHR expression reaches a functional threshold. This model is also widely assumed to apply to other bony vertebrates, including teleosts (Patiño et al., 2003).

Recently, we reported that the cloudy catshark (*Scyliorhinus torazame*) possesses a hierarchical ovary with strictly two follicles at each developmental stage (e.g. two F1, two F2, and so on), and lays exactly two eggs per reproductive cycle (Inoue et al., 2022). We further reported that all vitellogenic follicles abundantly express *lhr* (Arimura et al., 2024), suggesting that these follicles are capable of responding to the LH surge, as observed in birds. However, both ovulation and the robust P4 secretion driven by the upregulation of *steroidogenic acute regulatory protein 2* (*star2*) are restricted to F1 follicles following the LH surge in catshark (Inoue et al., 2026). This is in contrast to the avian system, in which both preovulatory and non-preovulatory follicles respond to the LH surge and stimulate P4 secretion. The cloudy catshark thus provides a unique model for dissecting the mechanisms that restrict ovulatory competence to F1 follicles within a hierarchical ovary.

In the present study, we sought to experimentally characterize the differences between F1 and F2 follicles by determining whether the LH surge drives distinct ovulation-associated responses. To address this, we performed RNA-seq analyses of F1 and F2 follicles before and after the endogenous LH surge and found that preovulatory F1 follicles exhibit qualitatively distinct and markedly stronger transcriptomic responses to the LH surge than F2 follicles. We then developed *ex vivo* follicle culture systems and demonstrated that an LH surge-like stimulation can induce robust P4 secretion and ovulation exclusively in F1 follicles. Although both F1 and F2 follicles were capable of receiving the LH signal, only F1 follicles activated ovulatory responses, including the pronounced P4 secretion accompanied by *star2* upregulation as well as the induction of ovulation-associated transcriptional programs. Notably, direct cAMP stimulation similarly induced P4 secretion in F1 but not F2 follicles, indicating that the differential ovulatory responsiveness of F1 and F2 follicles is determined downstream of cAMP signaling. These findings indicate that *lhr* expression and the capacity to receive the LH signal are not sufficient to confer ovulatory competence in this species. Instead, follicle-specific responsiveness to cAMP signaling downstream of LHR appears to determine ovulatory fate, challenging the prevailing assumption that follicle selection is determined by LHR abundance alone.

## 2. Materials and methods

### 2.1. Animals

Female cloudy catsharks (*Scyliorhinus torazame*) were transported from the Ibaraki Prefectural Oarai Aquarium to the Atmosphere and Ocean Research Institute (AORI) at The University of Tokyo. The animals were kept in a 1,000L experimental tank or 2,000L/3,000L holding tanks filled with recirculating natural seawater (36 ‰) maintained at 16 °C under a constant photoperiod (12 L: 12 D). They were fed with chopped sardines and squid to satiation, twice a week.

All experimental procedures were approved by the Animal Ethics Committee of Atmosphere and Ocean Research Institute of the University of Tokyo (P19-2). The present study was carried out in compliance with the ARRIVE guidelines.

### 2.2. Identification of reproductive phase by ultrasonography and plasma hormone monitoring

Ultrasonographic monitoring of the formation of encapsulated fertilized eggs was conducted daily underwater on unanesthetized females as described previously (Inoue et al., 2022). Meanwhile, the reproductive phase was determined by plasma hormone monitoring, distinguishing the pre-LH surge phase (T-phase) and the post-LH surge phase (P4-phase) (Inoue et al., 2026). Briefly, animals were anesthetized with 0.02 % (w/v) MS-222, and 0.1 mL of blood was collected every 12 h or 24 h from the caudal vasculature using a heparin-coated needle. Blood samples were centrifuged at 10,000 × *g* for 5 min to obtain the blood plasma, which was stored at -30 °C until hormone extraction. The plasma steroids were extracted and quantified by ELISA using commercially available kits (Cayman Chemicals, Ann Arbor, MI, USA) following the manufacturer’s protocols (Inoue et al., 2022). Individuals with plasma T concentrations higher than 5 ng/mL and plasma P4 concentrations lower than 5 ng/mL were classified as the “T-phase”, whereas individuals with plasma P4 concentrations higher than 5 ng/mL were categorized into the “P4-phase” group. The onset of P4 surge was operationally defined as the midpoint between the two consecutive sampling times when plasma P4 levels increased from less than 5 ng/mL to over 5 ng/mL. Sub-stages within the P4-phase were defined based on the time elapsed since the onset of P4 surge as follows.

Stage I, II, and III: Individuals sampled at 12 h, 36 h, and 42 h after the onset of the P4 surge were defined as Stages I, II, and III, respectively.

Stage IV: Individuals sampled ≥52 h after the onset of the P4 surge that exhibited ovulation shortly before sampling, with concurrent egg-capsule formation, were defined as Stage IV. For F1 follicles at Stage IV, postovulatory follicular tissue remaining in the ovary was sampled.

### 2.3. Tissue sampling and RNA extraction

Animals were terminally anesthetized with an overdose of MS-222 and euthanized by decapitation. For RNA extraction, egg yolk and oocytes were removed from the dissected ovarian follicles, and the remaining follicular layers were immediately frozen in liquid nitrogen in 2 mL screw-capped tubes containing 2.0φ zirconia beads (TOMY Seiko, Tokyo, Japan) and stored at -80 °C until use. Total RNA was extracted from the frozen tissues using ISOGEN (Nippon Gene, Tokyo, Japan), and treated with TURBO DNase (Thermo Fisher Scientific, Waltham, MA, USA) to remove genomic DNA, according to the manufacturers’ protocols.

### 2.4. RNA-sequencing

RNA concentration and RNA integrity number (RIN) of DNase-treated total RNA were measured using the QuantiFluor RNA System (Promega, Madison, WI, USA) and Agilent TapeStation system with RNA ScreenTape (Agilent Technology, Santa Clara, CA, USA), respectively (mean RIN ± SD: 7.1±1.2). For library preparation, 400 ng of total RNA was used to construct unstranded libraries using the NEBNext Ultra II RNA Library Prep Kit with Sample Purification Beads (New England BioLabs, Ipswich, MA, USA). Libraries with unique adapters were pooled and sequenced on an Illumina NovaSeq platform (Illumina, San Diego, CA) to generate 150-bp paired-end reads (200 Gb in total). Raw RNA-seq reads were trimmed for quality and adapter sequences using Trim Galore! v0.6.5, which incorporates Cutadapt v2.6 and FastQC v0.11.8, with default parameters (Martin, 2011). Trimmed reads were aligned to the catshark reference genome (Niwa et al., 2025) using STAR v2.7.11b (Dobin et al., 2013). These computational analyses were performed on the supercomputer at the ROIS National Institute of Genetics (Shizuoka, Japan). Gene-level read counts were obtained from genome-aligned BAM files using featureCounts (Liao et al., 2014). Global differences in gene expression profiles were assessed by calculating Bray–Curtis dissimilarities from TMM-normalized counts per million (CPM) values using the vegdist function in the vegan package (Oksanen et al., 2001). Principal coordinate analysis (PCoA) was performed by classical multidimensional scaling. Differential expression analyses were conducted separately for F1 and F2 follicles using DESeq2 (v1.40.2) (Love et al., 2014), and differentially expressed genes (DEGs) were defined as those with an adjusted p-value < 0.05 and an absolute log_2_ fold change > 1. Functional annotation was performed using eggNOG-mapper v2 (Cantalapiedra et al., 2021) and the KEGG database (Kanehisa et al., 2017). KEGG pathway enrichment analysis was conducted based on Fisher’s exact test, with multiple testing correction by the Benjamini–Hochberg method. Pathways with an adjusted p-value < 0.05 were considered significantly enriched. Cancer-related genes were defined as the combination of two gene sets: (1) genes annotated with KEGG pathway ko05200 (pathways in cancer) and (2) genes corresponding to KEGG orthologs assigned to ko05200 but not included in set (1). For genes not included in set (1), orthology was inferred by BLASTP searches against the NCBI RefSeq protein database of cloudy catshark with an E-value threshold of 5e-2, and the resulting hits were incorporated into set (2).

### 2.5. Preparation of ventral lobe extract (VLE)

For the reporter assay, ventral lobe extract (VLE) prepared by pooling the ventral lobes (VL) of the pituitary glands from 21 individuals in the previous study was used (Arimura et al., 2024). For follicle treatment, an additional 50 VLs were collected, and VLE was prepared following the same protocol and reconstituted in 8.3 mL of high-glucose DMEM (FUJIFILM Wako Pure Chemical, Osaka, Japan), with all reagent volumes proportionally scaled.

### 2.6. Recombinant LH

Recombinant cloudy catshark LH (rLH) and FSH (rFSH) was produced using Drosophila S2 cells transiently co-transfected with a pMT/BiP/V5-His A expression vector (Thermo Fisher Scientific, Waltham, MA, USA) encoding the *lhβ* or *fshβ* subunit linked to the g*lycoprotein hormone α (gpα)* subunit via a flexible peptide linker, together with the pCoBlast selection vector (Thermo Fisher Scientific, Waltham, MA, USA). Following transfection, cells were subjected to blasticidin selection, and protein expression was induced by CuSO₄. Cells were cultured under shaking conditions for 5 days following induction. The culture medium was collected and subjected to immobilized metal affinity chromatography using TALON cobalt resin (Takara Bio, Shiga, Japan). Bound proteins were eluted stepwise with elution buffers containing imidazole at concentrations of 5, 300, 400, and 500 mM. Eluted fractions were further purified by gel filtration chromatography using an ÄKTA FPLC system (Cytiva, Tokyo, Japan).

The presence and molecular weights of purified rLH and rFSH were confirmed by Western blot analysis. Proteins were separated by SDS–PAGE and transferred onto PVDF membranes, which were probed with a rabbit polyclonal anti-6×His tag primary antibody followed by an HRP-conjugated goat anti-rabbit IgG secondary antibody. Immunoreactive bands were detected using chemiluminescence, with expected molecular weights of approximately 27.6 kDa for rLH and 28.2 kDa for rFSH. Protein concentrations of the purified fractions were determined using a BCA protein assay. Purified rLH and rFSH fractions were either used directly for reporter assays (see below) or lyophilized, stored at -80 °C, and reconstituted in culture medium prior to follicular treatment.

### 2.7. Luciferase reporter assays

The reporter assay was conducted according to the methods described in our previous study (Arimura et al., 2024). In brief, HEK293A cells were maintained in high-glucose DMEM (FUJIFILM Wako Pure Chemical) supplemented with 10% fetal bovine serum and 1× penicillin–streptomycin at 37 °C in a 5% CO₂ incubator. The long-form cloudy catshark LH receptor (LHR-L; GenBank accession no. OR340974) was subcloned into the pcDNA3.1(+) expression vector, and was co-transfected with pGL4.29 CRE-luciferase vector and pGL4.74 (hRluc/TK) internal control vector (Promega) at a ratio of 4:1:1 to the cells using Lipofectamine 2000 (Thermo Fisher Scientific). After 6 h of transfection, the medium was replaced with 100 µL high-glucose DMEM containing the indicated hormones, and cells were incubated for an additional 11 h. Each treatment was performed in duplicate wells.

Following the incubation, the medium was removed and cells were lysed with 40 µL Passive Lysis Buffer (Promega). Cell lysates were stored at -80 °C until analysis. Luciferase activity was measured using the Dual-Luciferase Reporter Assay System (Promega). 20 µL of cell lysate was mixed with 35 µL Luciferase Assay Reagent II to measure firefly luciferase activity, followed by addition of 35 µL Stop & Glo Reagent to quench the firefly luminescence and measure Renilla luciferase activity. Dose–response curves were fitted using a four-parameter logistic model.

### 2.8. Follicle culture

A dissected ovary was placed in a clean petri dish containing catshark Ringer’s solution supplemented with antibiotics to remove blood and minimize microbial contamination. For whole follicle culture, F1 and F2 follicles were isolated and placed directly in a 12-well culture plate (IWAKI, Shizuoka, Japan). Follicles were pre-incubated for 3 h in 4 mL of M199-based medium (Supplementary Table 1) at 16 °C, and then transferred to a new 12-well plate with 4 mL of fresh culture medium containing VLE. Cultures were maintained at 16 °C for 72 h under ambient atmospheric conditions on a seesaw rocking platform (Nisshin Rika, Tokyo, Japan) at a speed of 16 tilts/minute. In vitro ovulation was monitored by continuous video recording using an HD webcam (Logitech, Lausanne, Switzerland).

For follicular layer strip culture, isolated follicles were longitudinally incised into eight equal strips along the animal–vegetal axis. The stripped follicular layers were washed in 1 mL of Ringer’s solution and transferred to a 48-well culture plate (IWAKI). Tissues were pre-incubated for 9 h in 200 µL of M199-based medium, and then transferred to a new 48-well plate containing 200 µL of culture medium for VLE, forskolin (Tokyo Chemical Industry, Tokyo, Japan) and dibutyryl-cAMP (Sigma-Aldrich, St Louis, MO, USA) treatment, or 140 µL of culture medium for recombinant LH treatment. For forskolin treatment, dimethyl sulfoxide (DMSO, Sigma-Aldrich) was used as the vehicle at a final concentration of 0.1% (v/v), while VLE, dibutyryl-cAMP and recombinant LH were dissolved directly in the culture medium. Tissues were incubated for up to 48 h under the same conditions as the whole follicle culture. After treatment, tissues and culture media were collected separately and stored at -80 °C and -30 °C, respectively, until further analysis. Steroid hormones were extracted from culture media and quantified following the procedures described previously (Inoue et al., 2026).

### 2.9. Reverse transcription quantitative PCR

500 ng RNA was reverse-transcribed into cDNA using the High Capacity cDNA Reverse Transcription Kit (Thermo Fisher Scientific). Quantitative PCR was performed using the KAPA SYBR FAST qPCR Kit (NIPPON Genetics, Tokyo, Japan) and gene-specific primers (Supplementary Table 2), as described previously (Inoue et al., 2026). Absolute transcript abundance was determined using standard curves generated from plasmids containing target sequences, and expression levels were normalized to β-actin as an endogenous control (Takagi et al., 2017).

### 2.10. Statistical analysis

Statistical analyses were performed using R (v4.5.1). Comparisons between measurements at a single concentration and the corresponding control from the same individuals were conducted using paired Student’s *t*-tests. Repeated-measures ANOVA followed by Tukey’s multiple comparison test was performed for dose-response experiments involving multiple concentrations. Time-course data were analyzed using linear mixed-effects models implemented in the lme4 package (v1.1.38), with individual identity included as a random effect to account for repeated measurements. Post hoc comparisons were performed using one-sided Dunnett’s tests, with comparisons made against the no-incubation group for gene expression analyses, or the 6 h post-VLE treatment group for P4-secretion.

## 3. Results

### 3.1. Dynamic transcriptomic transition in F1 follicles during the ovulatory process revealed by spatiotemporal RNA-seq

Within the single ovary of the cloudy catshark, follicles at multiple developmental stages exist simultaneously. The two F1 follicles are the largest and undergo ovulation following the LH surge, after which the two F2 follicles subsequently progress to become the next F1 follicles (Fig. 1a). A total of 22 F1 and 21 F2 samples were collected for RNA sequencing according to the time course shown in Fig. 1b. The T-phase samples correspond to the pre-LH surge stage, while four time points (stages I–IV) within the P4-phase represent post-LH surge stages (Fig. 1b). In F1 follicles, PERMANOVA revealed significant differences in transcriptomic composition between the T phase and each of the four P4-phase stages. Consistently, PCoA suggested a gradual shift in transcriptomic profiles with the progression of the stages; in particular, P4-phase stage IV samples occupied a distinct position relative to both the T phase and earlier P4-phase stages (Fig. 1c). In contrast, no significant differences were detected among T-phase and P4-phase stages in F2 follicles by PERMANOVA, and PCoA also showed no discernible stage-dependent shift in transcriptomic profiles (Fig. 1d). Although the comparison between the T-phase and P4-phase stage I in F2 follicles showed the lowest *P* value, it did not reach statistical significance (adjusted *P* = 0.06). When all F1 and F2 samples were analyzed together, a significant difference in transcriptomic profiles was observed between pooled F1 and F2 follicles. Notably, P4-phase stage IV F1 follicles were clearly separated from all other groups and differed markedly from P4-phase stage IV F2 follicles. In contrast, T-phase F1 and F2 follicles showed highly similar transcriptomic profiles (Supplementary Fig. 1a, b). Following ovulation of existing F1 follicles, F2 follicles at P4-phase stage IV

**Figure 1.**
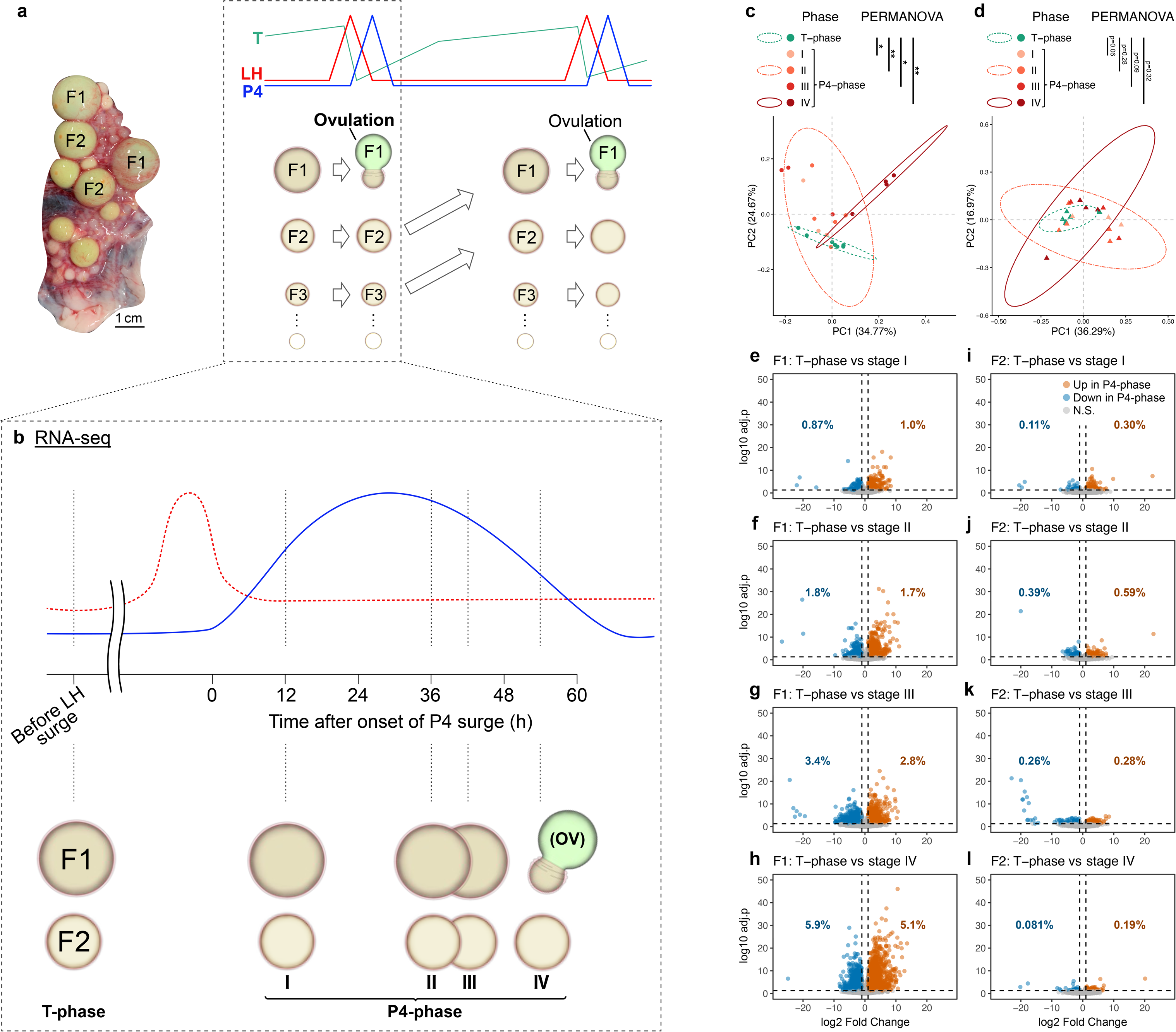
RNA-seq analysis reveals a transcriptomic shift in preovulatory F1 follicles, but not F2 follicles, following the endogenous LH surge a, Schematic representation of ovarian follicular hierarchy and associated hormonal dynamics. b, RNA-seq sampling design showing F1 (largest, ovulatory) and F2 (second-largest, non-ovulatory) follicles collected at the T-phase (pre-LH surge) and P4-phase (post-LH surge; stages I–IV). Stage IV corresponds to the post-ovulatory state of the F1 follicle. c, d, Principal coordinate analysis (PCoA) of F1 (c) and F2 (d) follicles based on Bray– Curtis distances. Separation between T- and P4-phase samples was assessed by PERMANOVA with 999 permutations. Ellipses represent 95% confidence intervals and are shown only for groups with n ≥ 4 biologically independent samples. \**P* < 0.05; \*\**P* < 0.01. e-l, Volcano plots showing differential gene expression between the T-phase and P4-phase stages I (e,i), II (f,j), III (g,k), and IV (h,l) in F1 (e–h) and F2 (i–l) follicles. Differentially expressed genes (DEGs) were defined as genes with adjusted *P* < 0.05 and |log_2_ fold change| > 1. The percentages of upregulated and downregulated genes in the P4-phase relative to the total number of genes are indicated in the upper right and upper left corners, respectively.

shifted toward the transcriptomic profile of T-phase F2 follicles—and, by extension, to T-phase F1 follicles—in the PCoA space, consistent with a progression toward the next follicular stage (Supplementary Fig. 1a, b). Consistent with these shifts, a large number of differentially expressed genes (DEGs) were identified between T-phase and P4-phase F1 follicles. In particular, approximately 11% of all genes exhibited differential expression between the F1 follicles of T-phase and P4-phase stage IV. In contrast, the number of DEGs in F2 follicles was more than three-fold lower, as illustrated by volcano plots (Fig. 1e-l).

### 3.2. Identification of F1 follicle–specific pathways enriched during the periovulatory P4 phase

To investigate pathways specifically associated with the ovulatory process in F1 follicles, KEGG pathway enrichment analyses were performed on DEGs between T-phase and P4-phase (Fig. 2). In F1 follicles, cancer-related pathways were the most frequently enriched across all comparisons between the T-phase and P4-phase, with all enrichments derived exclusively from genes upregulated at the P4-phase, as indicated in red color in the treemap plot (Fig. 2a, b and Supplementary Table S3). In contrast, cancer-related pathways were not enriched in F2 follicles, except for basal cell carcinoma, which was enriched among downregulated genes at P4-phase stage I (upper right in Fig. 2b and Supplementary Table S3). In F1 follicles at P4-phase stage IV, two cell growth-related pathways were enriched among downregulated genes, and a cell death-related pathway and two DNA replication/repair-related pathways were enriched among upregulated and downregulated genes, respectively, suggesting enhanced apoptotic activity (Fig. 2a). Among cancer-related genes, a total of 163 candidate target genes were identified whose expressions were specifically upregulated in the F1 follicles at P4-phase (stages I–IV) (Fig. 2c, Supplementary Fig. 2).

**Figure 2.**
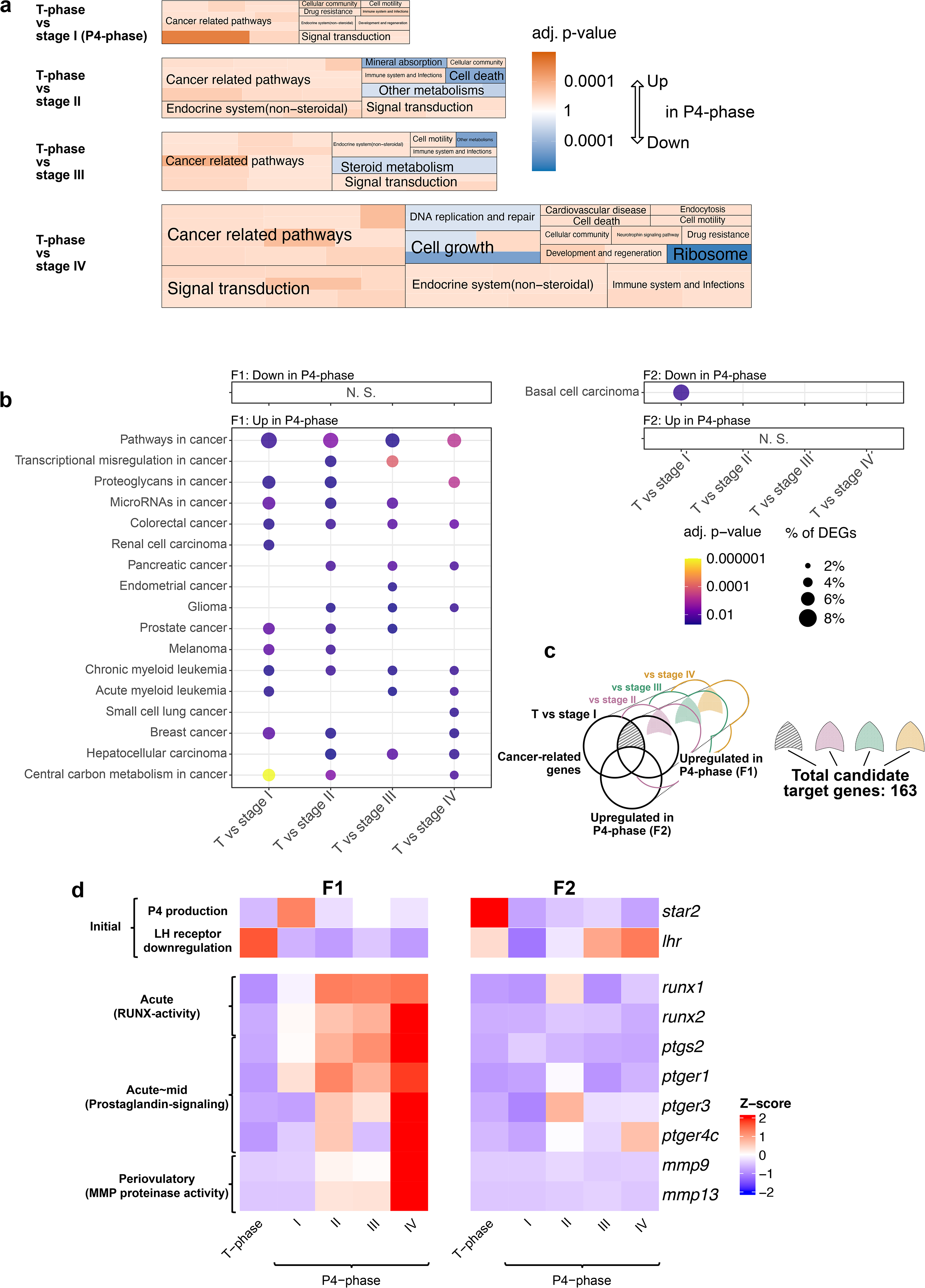
KEGG pathway analysis and target gene selection in preovulatory F1 follicles. a, Treemap visualization of KEGG pathway enrichment analyses comparing the T-phase with each P4-phase stage (I–IV). Enrichment analyses were performed separately for upregulated and downregulated DEGs. Color indicates adjusted *P* values (red, enriched among upregulated DEGs in P4-phase; blue, enriched among downregulated DEGs in P4-phase). Tile area represents the proportion of DEGs annotated to each pathway relative to the total number of upregulated or downregulated DEGs in each comparison. b, Bubble plot of enriched cancer-related pathways in F1 (left) and F2 (right) follicles. Bubble size represents the proportion of DEGs annotated to each pathway, and color indicates adjusted *P* values. c, Identification of candidate target genes defined as cancer-related genes upregulated in F1 follicles during the P4-phase (stages I–IV) relative to the T-phase, but not upregulated in F2 follicles at the corresponding stages. The union of stage-specific gene sets yielded a total of 163 genes. d, Heatmap showing z-score–normalized expression of representative candidate genes (*runx1, runx2, ptgs2, ptger1, ptger3, ptger4c, mmp9,* and *mmp13*) together with *star2* and *lhr*. Expression patterns of all 163 candidate genes are shown in Supplementary Fig. 2.

Prior to examining individual genes, we first examined the expression patterns of two genes previously implicated as being regulated by the LH surge: *star2*, which is upregulated in F1 follicles and downregulated in F2 follicles shortly after the LH surge (Inoue et al., 2026), and *lhr*, which is downregulated in both F1 and F2 follicles following the LH surge (Arimura et al., 2024). Consistent with the previous results, expression of *star2* was upregulated in F1 follicles and downregulated in F2 follicles at the P4-phase stage I in the RNA-seq data (Fig. 2d). Similarly, expression of *lhr* exhibited comparable downregulation at the P4-phase stage I of both F1 and F2 follicles in the current analyses. Notably, in F2 follicles, *lhr* expression showed a transient decrease followed by a recovery toward the later P4-phase (Fig. 2d).

To prioritize genes among the 163 candidate genes, we applied two criteria: (i) clear upregulation in F1 follicles, and (ii) prior evidence implicating a possible involvement in the ovulatory processes in vertebrates. Runt-related transcription factors (RUNX1 and RUNX2), both of which are implicated in ovulatory transcriptional regulation (Duffy et al., 2019; Lee-Thacker et al., 2020; Richards, 2007), were specifically upregulated in F1 follicles during the P4-phase, with *runx1* upregulated earlier than *runx2* (Fig. 2d). Genes involved in the prostaglandin synthesis and signaling were also identified. Prostaglandin-endoperoxide synthase 2 (PTGS2), a key enzyme in prostaglandin synthesis and essential for ovulation in both mammals and teleosts (Davis et al., 1999; Lister and Van Der Kraak, 2008; Needleman et al., 1986), showed F1-specific upregulation during the P4 phase. Prostaglandin receptor genes (*ptger1, ptger3*, and *ptger4c*) were also upregulated in periovulatory F1 follicles. Matrix metalloproteinases 9 and 13 (MMP9 and MMP13), LH-inducible proteases implicated in mammalian follicular rupture (Zhu, 2021), were upregulated in F1 follicles during the P4-phase, with particularly high expression at stage IV.

### 3.3. VLE treatment triggers F1-specific P4 secretion and ovulation

To determine whether LH stimulation induces follicle-specific responses, we examined the effects of pituitary ventral lobe extract (VLE) on isolated follicles under *ex vivo* culture conditions. VLE treatment induced ovulation in three out of four F1 follicles at 58, 59, and 61 h post-treatment (Fig. 3a, c; representative images at 72 h), whereas no ovulation was observed in any F2 follicles under the same conditions up to 72 h post-treatment (Fig. 3a). The remaining VLE-treated F1 follicle did not rupture; however, a pronounced protrusion was observed at the vegetal pole at 68 h post-treatment (Fig. 3b; representative images at 72 h). In follicles that underwent ovulation, the animal pole of the released oocyte faced the post-ovulatory follicle (POF) (Supplementary Fig. 3a, b), indicating that follicular rupture occurs on the opposite side of the oocyte, namely at the vegetal pole. Therefore, the follicle exhibiting a vegetal pole protrusion was interpreted as undergoing incomplete state of ovulation. Consistent with ovulatory competence, VLE treatment significantly increased P4 secretion in F1 follicles by ∼90-fold relative to saline control, whereas F2 follicles showed no detectable response. In parallel, *star2* expression showed an increase (∼10-fold) in F1 follicles following the VLE treatment, although this increase did not reach statistical significance (p=0.18).

**Figure 3.**
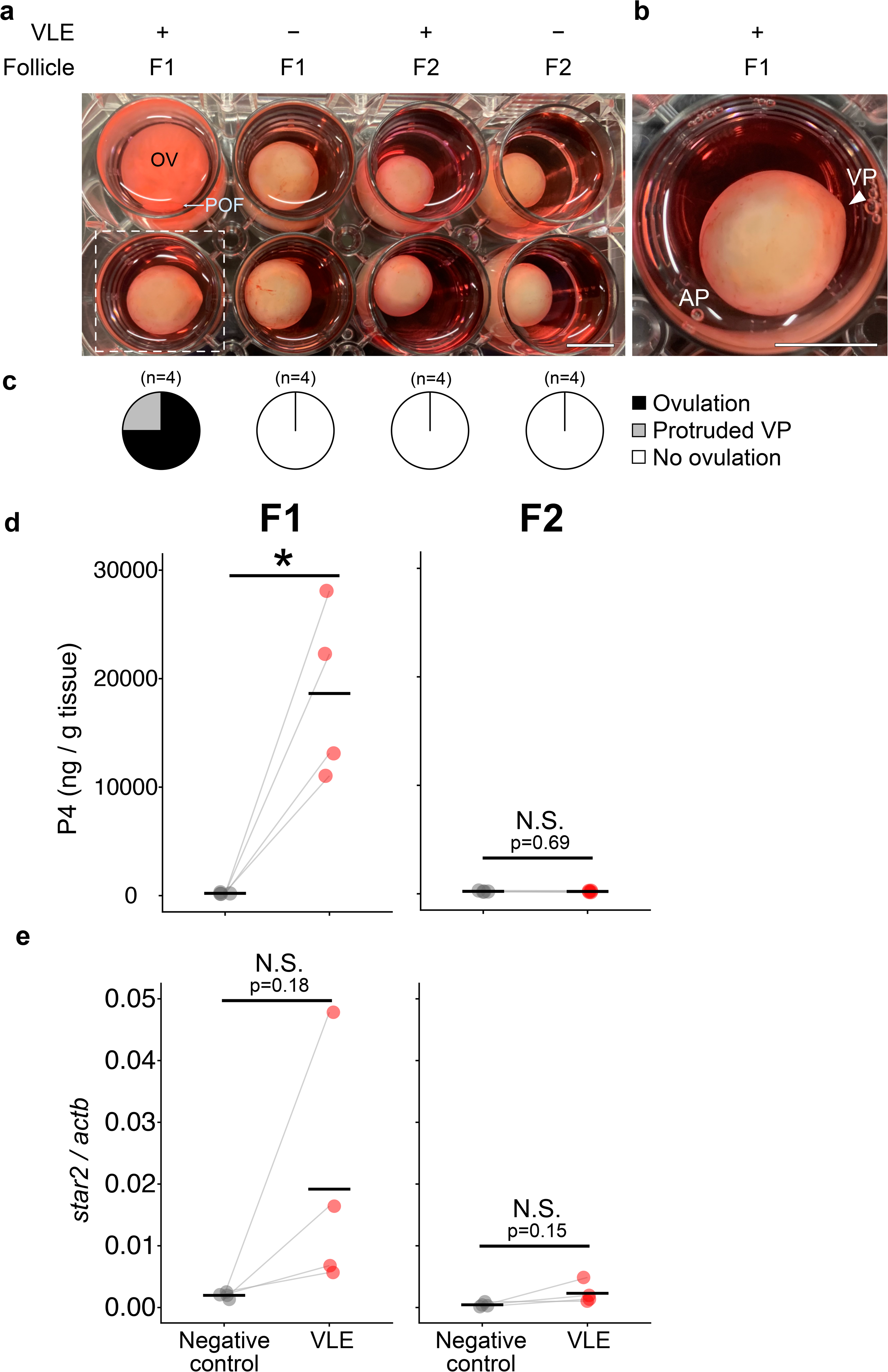
VLE treatment triggers ovulation and P4 secretion exclusively in F1 follicles. a, Representative images of F1 and F2 follicles from two individuals with or without VLE treatment. OV, ovum; POF, post-ovulatory follicular layer. The dashed outline indicates a VLE-treated F1 follicle that failed to ovulate and is shown at higher magnification in b. b, Higher-magnification view of a VLE-treated but unovulated F1 follicle. The vegetal pole (VP) shows outward protrusion. AP, animal pole; VP, vegetal pole. c, Ovulation rates in F1 and F2 follicles with or without VLE treatment. d, P4 secretion into the culture medium of F1 and F2 follicles with or without VLE treatment. e, Relative *star2* expression in F1 and F2 follicles with or without VLE treatment, normalized to *actb*. Statistical significance was determined by paired Student’s *t*-test. \**P* < 0.05. Scale bar, 1 cm.

### 3.4. F1 follicles require high concentration of LH to induce P4 secretion

To evaluate LH dose responsiveness, we performed luciferase reporter assays in HEK293 cells expressing the catshark LH receptor (Arimura et al., 2024). VLE induced dose-dependent activation of a cAMP response element (CRE) reporter in LHR-expressing cells (Fig. 4a). CRE activation was also detected in FSHR-expressing cells, but only at higher concentrations (≥5 µg/mL total protein) (Supplementary Fig. 4d), consistent with the presence of both LH- and FSH-producing cells in the pituitary ventral lobe (Arimura et al., 2024; Quérat et al., 2001). To test whether effects of VLE are mediated by LH, we generated recombinant catshark LH (rLH) (Supplementary Fig. 4). Immunoreactive bands of the expected molecular size were detected in purified fractions 12 and 13 following high-performance liquid chromatography (Supplementary Fig. 4b, c), confirming successful expression and purification. rLH induced dose-dependent CRE activation, with dose–response curves parallel to those of VLE (Fig. 4a). The EC₅₀ values were 0.064 µg/mL for VLE and 0.0085 µg/mL for rLH.

**Figure 4.**
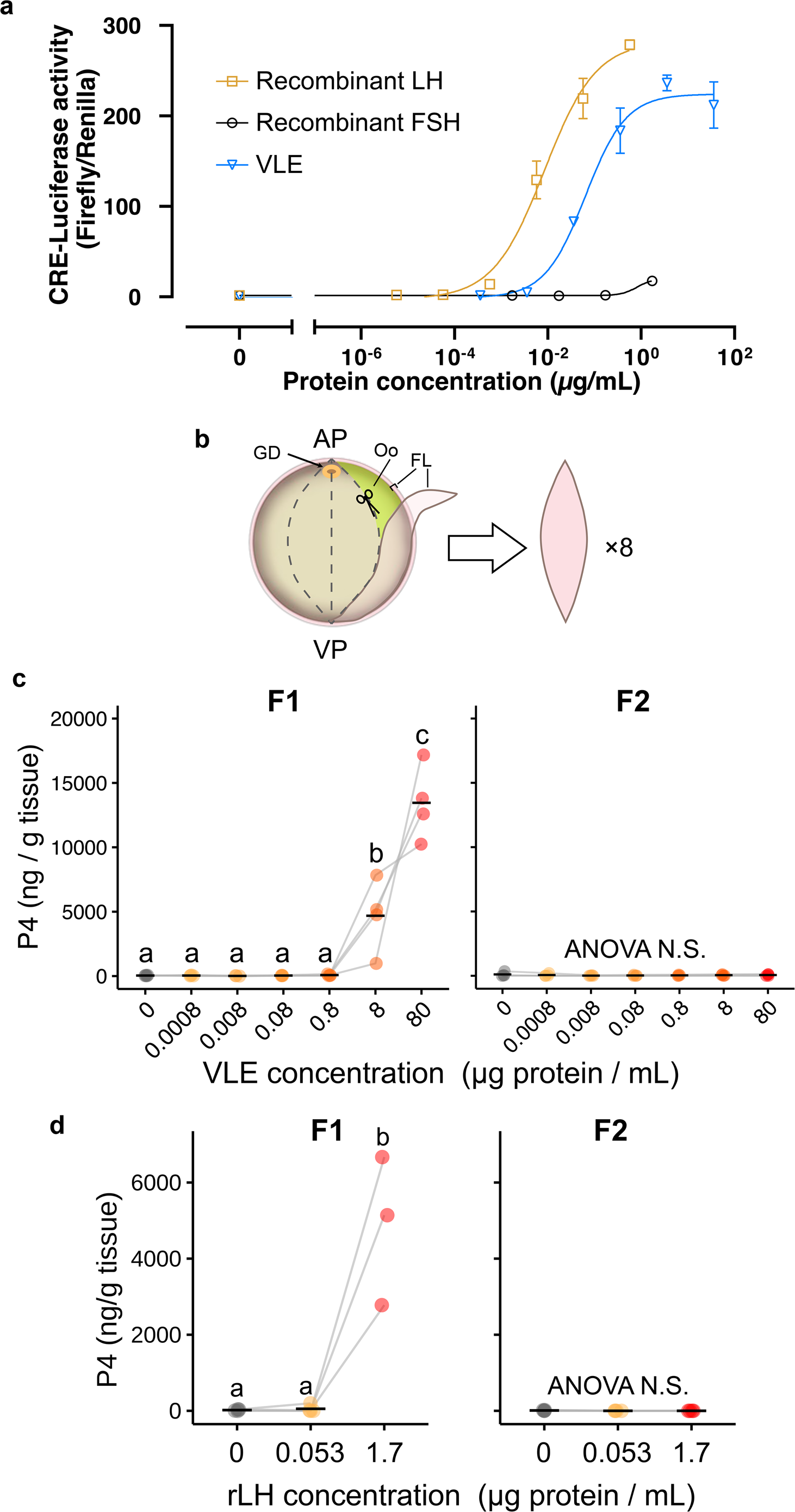
Dose-dependent effects of VLE and recombinant LH on LHR activation and P4 secretion in F1 and F2 follicles. a, Reporter assay of catshark LHR (long isoform) activity in HEK293A cells using recombinant LH (rLH), recombinant FSH (rFSH), or VLE. The y-axis represents relative CRE-driven firefly luciferase activity normalized to Renilla luciferase. The x-axis indicates the total protein concentration of the recombinant hormones or VLE. b, Schematic diagram of follicular layer strip preparation. The follicular layer (FL) was longitudinally divided into eight equal strips along the animal-vegetal axis. AP, animal pole; VP, vegetal pole; Oo, oocyte; FL, follicular layer; GD, germinal disk. c, P4 secretion from F1 and F2 follicular layer strips treated with increasing concentrations of VLE. d, P4 secretion from F1 and F2 follicular layer strips treated with rLH. Statistical significance was determined by repeated-measures ANOVA followed by Tukey’s multiple comparison test. \**P* < 0.05.

To further investigate dose-dependent effects of VLE treatment, we newly established a “ follicular layer strip culture” system as illustrated in Fig. 4b. Each follicle was longitudinally sectioned into eight equal strips along the animal–vegetal axis (Fig. 4b), based on the observation that follicular rupture occurs at the vegetal pole (Fig. 3b; Supplementary Fig. 3) and the assumption of axial symmetry along this axis. This approach enabled us to perform parallel testing of multiple conditions using a single follicle. In this system, VLE treatment induced P4 secretion in F1 follicles only at concentrations above 8 µg/mL, with a pronounced increase at 80 µg/mL, whereas F2 follicles remained unresponsive even at the highest doses tested (Fig. 4c). Similarly, the rLH induced P4 secretion exclusively in F1 follicles at the highest concentration tested (1.7 µg/mL; Fig. 4d), with no detectable response in F2 follicles.

### 3.5. VLE treatment induces temporally ordered transcriptional activation in F1 follicles

We next investigated whether candidate target genes identified by RNA-seq are directly responsive to LH stimulation in F1 follicles. Following the 48 h treatment with a high dose of VLE (80 µg/mL), the expression of *lhr* significantly decreased in both F1 and F2 follicles (Fig. 5a). VLE treatment coordinately increased expressions of *star2, runx1, runx2, ptgs2, ptger1*, and *mmp9* in F1 follicles, whereas induction was absent or minimal in F2 follicles (Fig. 5b–g). Although *mmp13* expression did not reach statistical significance in F1 follicles (p = 0.07), a marked increase (∼23-fold compared to control) was observed in the VLE-treated group, suggesting a strong but variable response. In contrast, F2 follicles showed only a modest (∼1.9-fold) but statistically significant induction. Meanwhile, other prostaglandin receptor paralogs (*ptger3* and *ptger4c*) exhibited only modest or non-significant changes in both F1 and F2 follicles (Supplementary Fig. 7).

**Figure 5.**
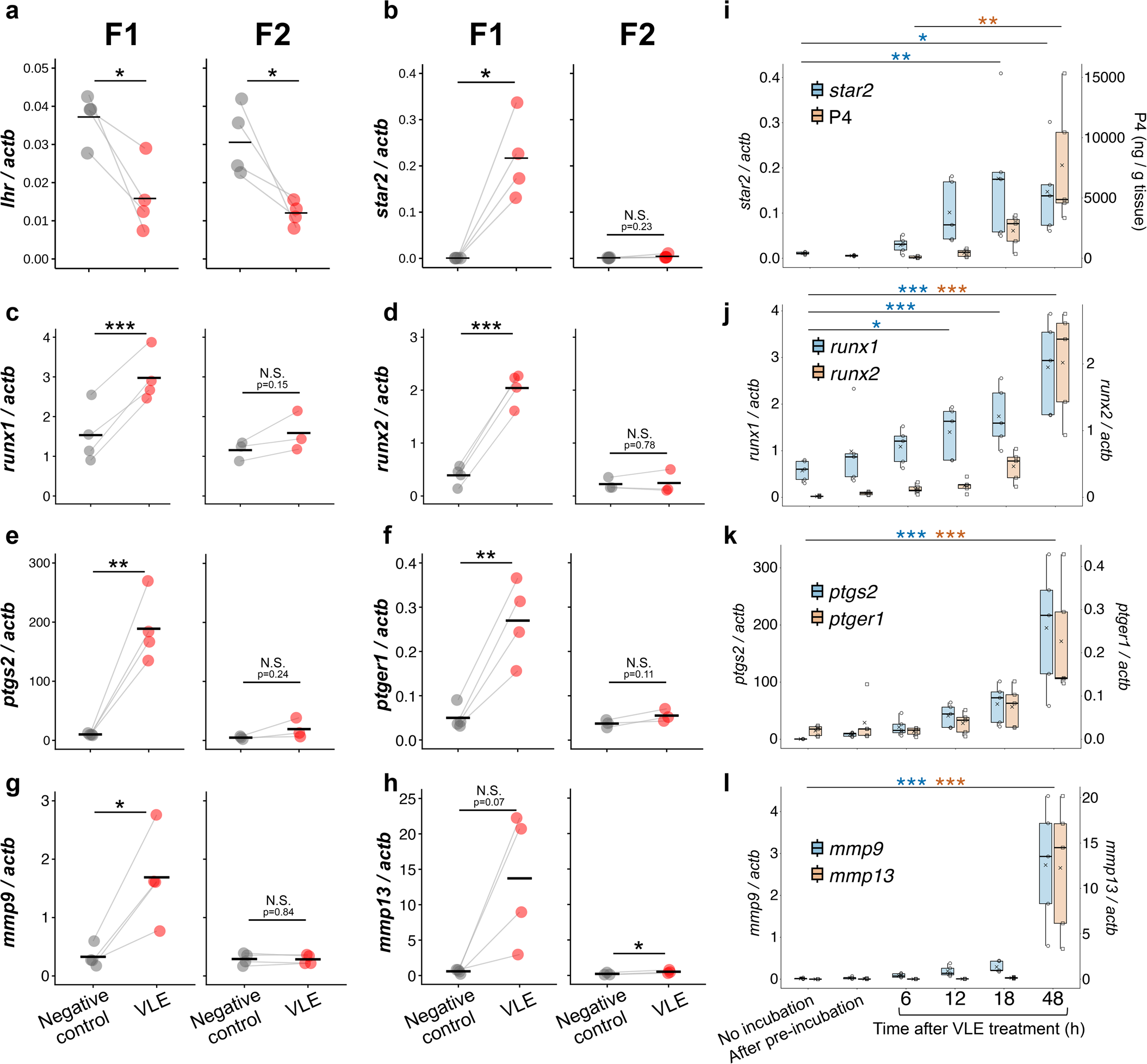
VLE treatment induces time-dependent ovulation-related gene expression and P4 secretion specifically in F1 follicles. a–h, Relative expression of *lhr*, *star2*, *runx1*, *runx2*, *ptgs2*, *ptger1*, *mmp9,* and *mmp13* in F1 (left) and F2 (right) follicles treated with 0 or 80 µg ml⁻¹ VLE (total protein). i–l, Time course of ovulation-related gene expression and P4 secretion in VLE (80 µg ml⁻¹)-treated F1 follicles. Samples were collected at no incubation, after pre-incubation, and 6, 12, 18 and 48 h after VLE treatment. i, *star2* expression and P4 secretion; j, *runx1* and *runx2* expression; k, *ptgs2* and *ptger1* expression; l, *mmp9* and *mmp13* expression. Gene expression levels were normalized to *actb*. For a–h, paired Student’s *t*-tests were used to compare VLE-treated and control groups. Time-course data (i–l) were analyzed using linear mixed-effects models. Post hoc comparisons against the no-incubation group (gene expression) or the 6 h post-VLE group (P4 secretion) were performed using one-sided Dunnett’s tests. \**P* < 0.05; \*\**P* < 0.01; \*\*\**P* < 0.001.

To further characterize the temporal dynamics of LH-responsive genes, a time-course analysis was performed in F1 follicles. VLE treatment increased the expression of seven genes in F1 follicles, namely *star2, runx1, runx2, ptgs2, ptger1*, *mmp9,* and *mmp13* in a temporally coordinated manner. The increase in *star2* expression was observed 12 h after treatment and reached statistical significance by 18 h. The release of P4 increased slightly following the upregulation of *star2* expression, and became significant at 48 h after VLE treatment (Fig. 5i). The expression of *runx1* was relatively high even before VLE treatment, increased significantly at 12 h after VLE treatment, and continued to increase until 48 h. On the other hand, expression of *runx2* showed a delayed response, reaching significant upregulation at 48 h, with a modest increase at 18 h (Fig. 5j). The expressions of *ptgs2* and *ptger1* exhibited similar temporal patterns to *runx2*, suggesting their involvement during the mid-phase ovulatory processes (Fig. 5k). Finally, *mmp9* and *mmp13* were drastically upregulated at 48 h, with fold changes of approximately 9 and 90, respectively, relative to 18 h (Fig. 5l). These temporal expression patterns closely mirror those observed in the RNA-seq analyses.

### 3.6. cAMP stimulation induces P4 secretion in F1 but not F2 follicles

LHR is a Gs-coupled GPCR, and LH stimulation increases intracellular cAMP levels, thereby regulating downstream transcription (Fig. 4a). To determine whether F2 follicles retain the ability to respond to cAMP, F1 and F2 follicles were treated with forskolin, which directly increases intracellular cAMP levels independently of GPCR activation. P4 secretion was measured as a functional indicator of the ovulatory responsiveness. Forskolin treatment significantly increased P4 secretion exclusively in F1 follicles, resulting in an approximately 60-fold increase relative to the control (Fig. 6a). Furthermore, treatment with dibutyryl-cAMP, a membrane-permeable cAMP analogue, similarly induced P4 secretion in F1 follicles at 1 mM (25-fold increase relative to control), whereas no significant increase was observed in F2 follicles (Fig. 6b).

**Figure 6.**
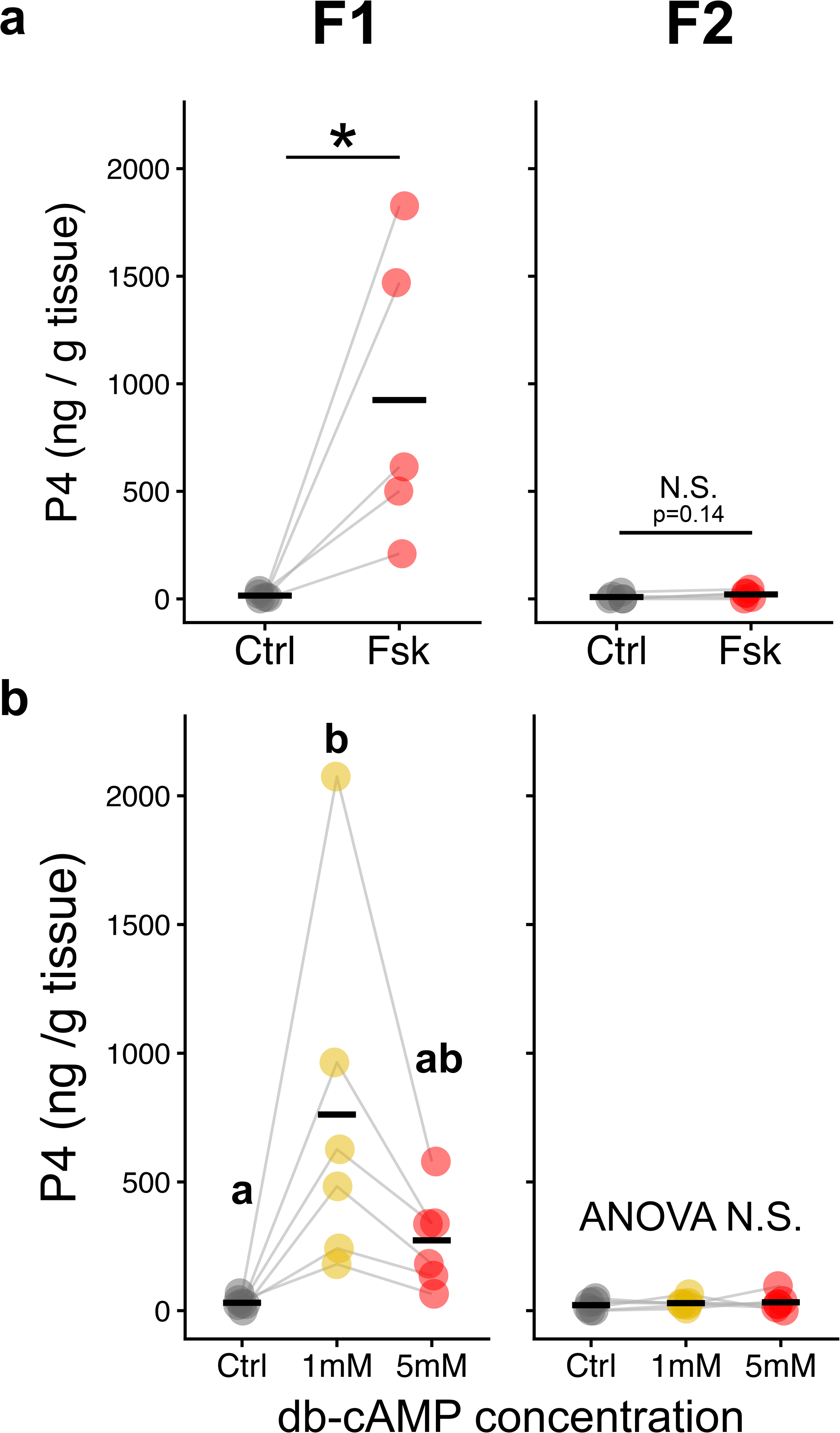
Direct cAMP stimulation specifically triggers P4 secretion from F1 follicles. a, P4 secretion from F1 and F2 follicular layer strips treated with 0.1% DMSO (vehicle control) or forskolin (10 µM). b, P4 secretion from F1 and F2 follicular layer strips treated with medium alone (control) or dibutyryl-cAMP (1 mM or 5 mM). Statistical significance was determined by paired Student’s *t*-test (a) or repeated-measures ANOVA followed by Tukey’s multiple comparison test (b). \**P* < 0.05.

## 4. Discussion

The present study demonstrates that preovulatory F1, but not F2, follicles in the cloudy catshark exhibit ovulatory responses to the LH surge. Our previous studies demonstrated comparable levels of *lhr* expression in F1 and F2 follicles, as well as decreases in *lhr* expression and T secretion in both follicle classes following the LH surge (Arimura et al., 2024; Inoue et al., 2026), indicating that both F1 and F2 follicles are capable of receiving and responding to the LH signal. Downregulation of *lhr* following LH stimulation was also confirmed in both F1 and F2 follicles using the *ex vivo* culture system (Figs. 2d and 5a). However, LH stimulation selectively activated ovulation-associated transcriptional programs, including pathways overlapping with cancer-associated processes, as well as *star2*-dependent P4 synthesis, exclusively in F1 follicles (Fig. 2). These responses led to P4 secretion and ovulation in F1 follicles but not in F2 follicles (Figs. 2, 3 and 5). Furthermore, direct activation of the cAMP pathway, which bypasses LHR activation, induced P4 production only in F1 follicles, indicating that the critical difference in ovulatory responsiveness between F1 and F2 follicles lies downstream of cAMP (Fig. 6). These findings differ from the prevailing model in several other vertebrates, in which differential LHR expression and LH responsiveness are closely associated with follicular ovulatory competence. Instead, our results indicate that follicular ovulatory fate is determined, at least in part, by differential responsiveness downstream of cAMP rather than by LHR abundance.

### Ovulatory pathways in F1 follicles

The transition of F1 follicles from the T-phase to the ovulatory P4-phase was accompanied by pronounced transcriptomic changes compared with F2 follicles (Fig. 1c, e–h; Supplementary Table 3). Notably, cancer-related pathways were prominently enriched (Fig. 2a,b), and this is in line with the concept that ovulation represents an inflammatory tissue-remodeling process, which shares transcriptional pathways with tumorigenesis (Richards, 2018). In mammals, RUNX1 and RUNX2 are known to play key roles in ovulatory processes (Dinh et al., 2023; Liu et al., 2009). LH-induced RUNX1 promotes *runx2* expression, and RUNX1 and RUNX2 cooperatively promote *mmp13* expression, potentially facilitating follicular rupture (Park et al., 2010). In catshark, we found that *runx1* was upregulated more rapidly than *runx2* and *mmps* following VLE treatment (Fig. 5j-l). Likewise in mammals, this LH-induced RUNX1 may bind to the highly conserved RUNX-binding motifs (5’-TGTGGT-3’) identified in *runx2* and *mmp13* (Supplementary Fig. 8), thereby inducing the subsequent upregulation of *runx2* and *mmps* following LH stimulation.

A second notable cascade activated in ovulatory F1 follicles is the prostaglandin signaling. LH-dependent prostaglandin synthesis via PTGS2, an enzyme that synthesizes the prostaglandin precursor (Needleman et al., 1986), is indispensable for ovulation in both mammals and teleosts (Davis et al., 1999; Ogiwara et al., 2023; Tang et al., 2017). In mammals, prostaglandins produced in follicular cells primarily act through PTGER2 and PTGER4 in cumulus cells during ovulation (Niringiyumukiza et al., 2018), whereas in teleosts they signal via PTGER4b expressed in granulosa cells (Takahashi et al., 2018). Our results demonstrate that both prostaglandin synthesis (*ptgs2*) and signaling (*ptger1*) are specifically induced by LH in F1 follicles (Fig. 5e, f, k). These results suggest a conserved requirement for prostaglandins in vertebrate ovulation, accompanied by potential diversification in receptor usage.

*mmp9* and *mmp13* were upregulated in periovulatory F1 follicles (Figs. 2d and 5l). LH-induced MMP9 is reported to be associated with follicular rupture in a wide range of vertebrates (Hrabia et al., 2019; Kim and Yoon, 2020; Liu et al., 2018). Also in tunicates, an ancestral *mmp2/9/13* gene is essential for ovulation (Matsubara et al., 2019). These MMPs are thought to contribute to ovulation by degrading the follicular basal lamina and extracellular collagens (Zhu, 2021). On the other hand, while these genes are upregulated in response to LH, studies in mouse and zebrafish suggest that they are not directly required for ovulation (Itoh et al., 1999; Silva et al., 2020), but may instead contribute to the regression of post-ovulatory follicular layers or corpora lutea (Levin et al., 2018). In catshark, periovulatory activation of MMP9 and MMP13 may facilitate follicular rupture through degradation of the follicular wall (Fig. 2d; Fig. 5l). However, definitive determination of their roles in ovulation will require functional validation using loss-of-function approaches.

### What is the difference between F1 and F2?

The present findings demonstrate that LH signaling induces P4 secretion and ovulation exclusively in F1 follicles. What, then, distinguishes ovulatory F1 from non-ovulatory F2 follicles, and accounts for their differential activation of ovulation-associated transcriptional programs? One clear conclusion is that this distinction cannot simply be attributed to differences in LH receptivity. Consistent with our previous findings (Arimura et al., 2024), *lhr* is expressed at comparable levels in F1 and F2 follicles during the T-phase (Figs. 2d and 5a). The present *ex vivo* culture experiments further demonstrated that VLE treatment downregulated *lhr* expression in both F1 and F2 follicles, consistent with our previous observation that *lhr* expression markedly decreased in both follicle classes during the P4-phase following the LH surge (Arimura et al., 2024). Given that LH-induced downregulation of LHR has been shown to occur through cAMP-dependent pathways (Gulappa et al., 2015), the observed downregulation of *lhr* in both F1 and F2 follicles may reflect an increase in intracellular cAMP following LH stimulation (Fig. 5a). Indeed, the reporter assay using catshark LHR suggested that LH stimulation induced cAMP production (Fig. 4a), and P4 production in F1 follicles was directly induced by forskolin and dibutyryl-cAMP treatments (Fig. 6). Nevertheless, F2 follicles failed to secrete P4 in response to direct activation of the cAMP pathway (Fig. 6). If the lack of P4 secretion in F2 follicles were simply due to the absence of LHR or an inability to receive the LH signal, direct activation of the cAMP pathway, thereby bypassing LHR activation would be expected to induce P4 secretion; however, it did not. These results indicate that the critical determinant of ovulatory competence lies downstream of cAMP, and that only F1 follicles are capable of translating the cAMP signal into ovulation-associated transcriptional activation, whereas F2 follicles may lack the molecular context required for this response. Future studies should therefore seek to identify the factors downstream of cAMP that distinguish F1 from F2 follicles, including differences in the permissiveness of the epigenetic landscape and the availability of specific transcriptional cofactors.

It is also noteworthy that F1 and F2 follicles exhibit distinct patterns of change as the egg-laying cycle progresses. During the periovulatory period, F1 follicles underwent the most pronounced transcriptional changes, as reflected by the highest proportion of DEGs and their distinct clustering in the PCoA at P4-phase stage IV (Fig. 1c, h; Supplementary Fig. 1a, b). Following ovulation, F1 follicles activated apoptotic pathways (Fig. 2a), consistent with the previously reported regression of post-ovulatory follicular layers in this species (Arimura et al., 2024). By contrast, in F2 follicles, the LH-induced decrease in *lhr* expression was restored to pre-stimulation levels following F1 ovulation, indicating recovery of their capacity to respond to LH (Fig. 2d). Moreover, F2 follicles returned to a T-phase-like transcriptional profile following F1 ovulation (Supplementary Fig. 1). As the follicular hierarchy advances, these F2 follicles subsequently transition to F1 position and must therefore acquire the capacity to respond to cAMP, and to initiate P4 production and ovulation following the next LH surge. Thus, the transition from F2 to F1 appears to involve the acquisition of cAMP responsiveness and, consequently, ovulatory competence. Elucidating the mechanisms underlying this transition within the follicular hierarchy will be an important subject for future investigation.

### Comparative perspectives of differential LH receptivity and ovulation competence in vertebrates

The concept that differential responsiveness to LH surge underlies variation in ovulation number is widely accepted across vertebrates. Ovulation number has commonly been interpreted within an LHR-threshold model, where the number of follicles expressing LHR above a critical level determines how many follicles proceed to ovulation.

In mammals, only the dominant follicle(s) acquires sufficient levels of LHR to respond fully to the LH surge and proceeds to ovulation, whereas the remaining follicles undergo atresia (Shilo et al., 2022). Accordingly, the pool of mature follicles capable of responding to the LH surge is generally thought to correspond to the population destined to ovulate (Fig. 7; Longo et al., 2025; Zeleznik, 2004). Notably, direct stimulation with forskolin or a cAMP analogue can induce P4 secretion even in immature, non-preovulatory follicles (Hedin and Rosberg, 1983), suggesting that the downstream cAMP-responsive machinery is already functional before follicles acquire full ovulatory competence. These observations are consistent with a model in which acquisition of sufficient LHR expression, rather than competence of the downstream cAMP pathway itself, serves as a major gate for ovulatory fate.

**Figure 7.**
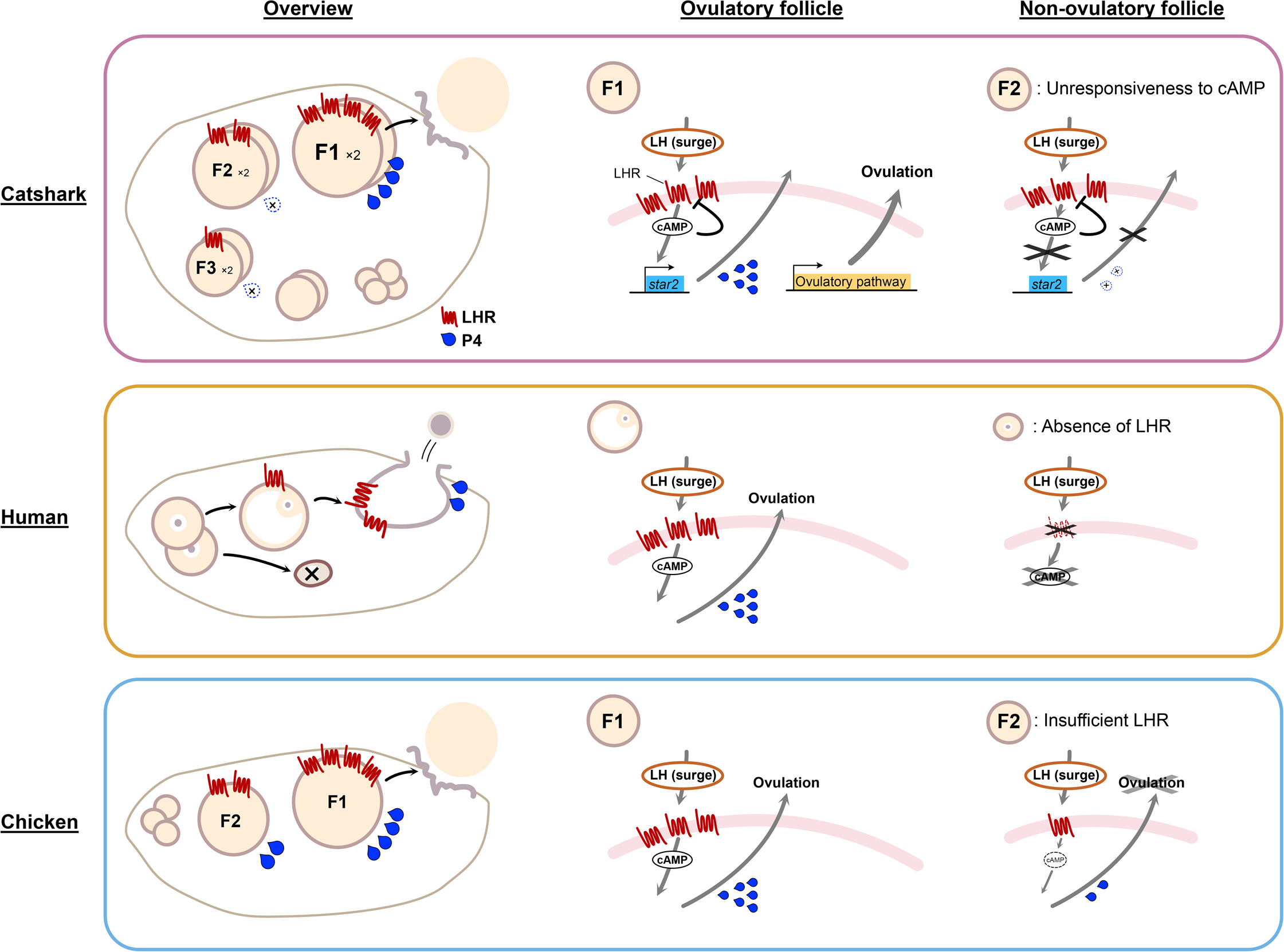
Schematic representation of diverse mechanisms of ovulatory follicle selection in vertebrates. Comparison of follicle development, selection, and ovulation mechanisms. In human, a subset of primordial follicles is recruited for growth, while the majority remain arrested. Among growing follicles, only the selected dominant follicle acquires LHR expression in granulosa cells and proceeds to ovulation in response to the LH surge. In chicken, follicles are organized into a hierarchical structure. LHR expression increases progressively during vitellogenic growth (F3 to F1 in this figure). Both preovulatory F1 and non-preovulatory follicles (e.g., F2 and earlier stages) respond to the LH surge by secreting progesterone (P4), although the response is markedly stronger in F1 follicles, which exhibit higher LHR expression. In catshark, follicles also form a hierarchy (F3 to F1 in this figure) with progressively increasing LHR expression, similar to chicken. However, despite this comparable ability to perceive the LH surge, only the pair of F1 follicles produce P4 and activate ovulation-associated gene expression, thereby restricting ovulation to the F1 follicle.

In birds, as in the catshark, follicles develop simultaneously in a hierarchical ovary, with one F1, one F2, and so on present at a given time. Both F1 and F2 follicles, and occasionally follicles at earlier-stages, express LHR (Fig. 7; Johnson et al., 1996) and exhibit qualitatively similar responses to the LH surge, including P4 secretion (Johnson et al., 2002) and protease expression (Jackson et al., 1993). However, ovulation occurs exclusively in the F1 follicle, which exhibits the highest *lhr* expression and quantitatively greater responses to LH stimulation (Fig. 7). Similar to mammals, non-ovulatory follicles in birds can also produce P4 in response to direct cAMP stimulation, in clear contrast to our findings in catshark (Dean et al., 1995; Sechman et al., 2011). Accordingly, ovulation in birds is thought to occur when LHR levels surpass a functional threshold. Thus, variation in the number of follicles ovulated in response to a single LH surge has traditionally been attributed to variation in the number of follicles expressing LHR above the threshold required for ovulatory competence (Liu et al., 2022b; Zeleznik, 2004).

By contrast, our findings indicate that ovulation number in the catshark is determined not simply by differential LH receptivity, but by differential activation of ovulation-associated responses downstream of cAMP. In the catshark, both F1 and F2 follicles express *lhr* (Fig. 2d, Fig. 7; Arimura et al., 2024) and appear capable responding to the LH surge. This is supported by *lhr* downregulation following LH stimulation in both F1 and F2 follicles (Fig. 5a), a response also observed in birds and mammals (Kishi et al., 2018; Yamamura et al., 2001). Nevertheless, ovulation-associated responses, including P4 secretion and the activation of protease-related transcriptional pathways, were induced in F1 but not F2 follicles (Fig. 7). We previously reported that the cloudy catshark ovulates and lays only two eggs per reproductive cycle (Inoue et al., 2022). The uncoupling of LH receptivity from downstream cAMP responsiveness may therefore provide a mechanistic explanation for the strict limitation in ovulation number in this species, representing a regulatory feature that appears to distinguish the catshark from previously characterized vertebrate models. Overall, the cloudy catshark provides a unique model for studying follicular selection and offers new perspectives on the mechanisms that determine ovulatory fate beyond the conventional model for ovulatory regulation.

### Future perspectives

Building on the first functional demonstration that LH serves as an essential endocrine trigger for ovulation in cartilaginous fishes, the experimental platforms established in this study provide new opportunities to investigate ovulatory mechanisms in this lineage, with important implications for understanding functional evolution across vertebrates. The newly developed follicular layer strip culture system, combined with VLE and rLH, enables mechanistic investigations using a limited number of follicles and individuals. Application of this system is expected to facilitate elucidation of why only F1 follicles respond to the LH surge and proceed to ovulation in the cloudy catshark, which possesses a hierarchical ovary. Furthermore, these approaches will contribute to a broader understanding of the common mechanisms by which gonadotropins regulate follicular maturation and ovulation in vertebrates. Taken together, the current methodological advances provide a foundation for future functional investigations and further establish the cloudy catshark as a powerful emerging model for dissecting hierarchical follicle selection and the evolution of vertebrate reproductive strategies.

## CRediT authorship contribution statement

**Ryotaro Inoue:** Conceptualization, Data curation, Formal analysis, Funding acquisition, Investigation, Methodology, Project administration, Resources, Validation, Visualization, Writing – original draft, Writing – review & editing.

**Takanori Kinugasa:** Data curation, Investigation, Methodology, Validation, Writing – review & editing.

**Keigo Nagasaka:** Data curation, Resources, Validation, Writing – review & editing.

**Kotaro Tokunaga:** Resources, Writing – review & editing.

**Shigeho Ijiri:** Conceptualization, Methodology, Funding acquisition, Project administration, Resources, Supervision, Validation, Writing – review & editing.

**Susumu Hyodo:** Conceptualization, Funding acquisition, Project administration, Resources, Supervision, Validation, Visualization, Writing – original draft, Writing – review & editing.

## Data availability

Data will be made available on request.

## Acknowledgements

We thank Dr. Marty Kwok Shing Wong for valuable discussions and comments on the manuscript. We also thank Mr. Hikaru Ishihara and Dr. Shinji Kanda for their assistance with RNA-seq library preparation and discussion. We are grateful to Drs. Wataru Takagi, Yung-Che Tseng and Ling Chiu for valuable advice and comments on RNA-seq analysis.

## Ethical approval

All animal experiments were conducted according to the Guidelines for Care and Use of Animals approved by the committees of The University of Tokyo (P19-2). The present study was carried out in compliance with the ARRIVE guidelines.

Consent to participate/consent to publish Not applicable.

## Abbreviations

cAMP: cyclic adenosine monophosphate;
LH: Luteinizing hormone;
*lhr*: *Luteinizing hormone receptor*;
*mmp9*: *Matrix metalloproteinase 9*;
mmp13: *Matrix metalloproteinase 13*;
POF: Post ovulatory follicular layer;
P4: Progesterone;
*ptgs2*: *Prostaglandin-endoperoxide synthase 2*;
*ptger1*: *Prostaglandin E receptor 1*;
*ptger3*: *Prostaglandin E receptor 3*;
*ptger4c*: *Prostaglandin E receptor 4c*;
rLH: Recombinant luteinizing hormone;
*runx1*: *Runt-related transcription factor 1*;
*runx2*: *Runt-related transcription factor 2*;
*star2*: *Steroidogenic acute regulatory protein 2*;
T: Testosterone;
VLE: Ventral lobe extract;

---

**Supplementary Figure 1.**
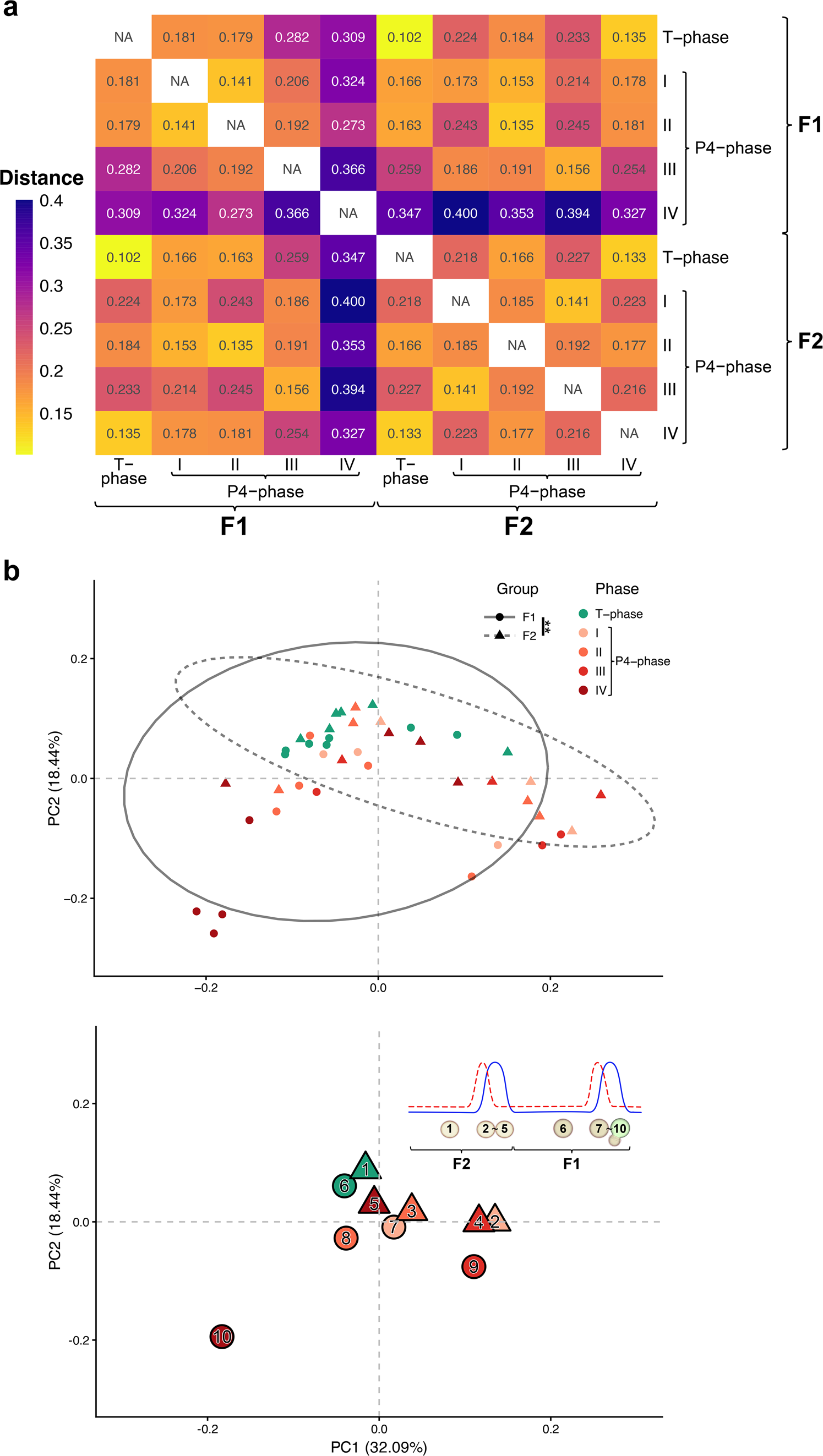
Bray–Curtis distance and principal coordinate analysis (PCoA) of pooled F1 and F2 follicles, related to Figure 1. a, Bray–Curtis distance matrix of pooled F1 and F2 follicles. b, Principal coordinate analysis (PCoA) based on Bray–Curtis distances. Separation between F1 and F2 samples was assessed by PERMANOVA with 999 permutations. Ellipses represent 95% confidence intervals. \*\**P* < 0.01.

**Supplementary Figure 2.**
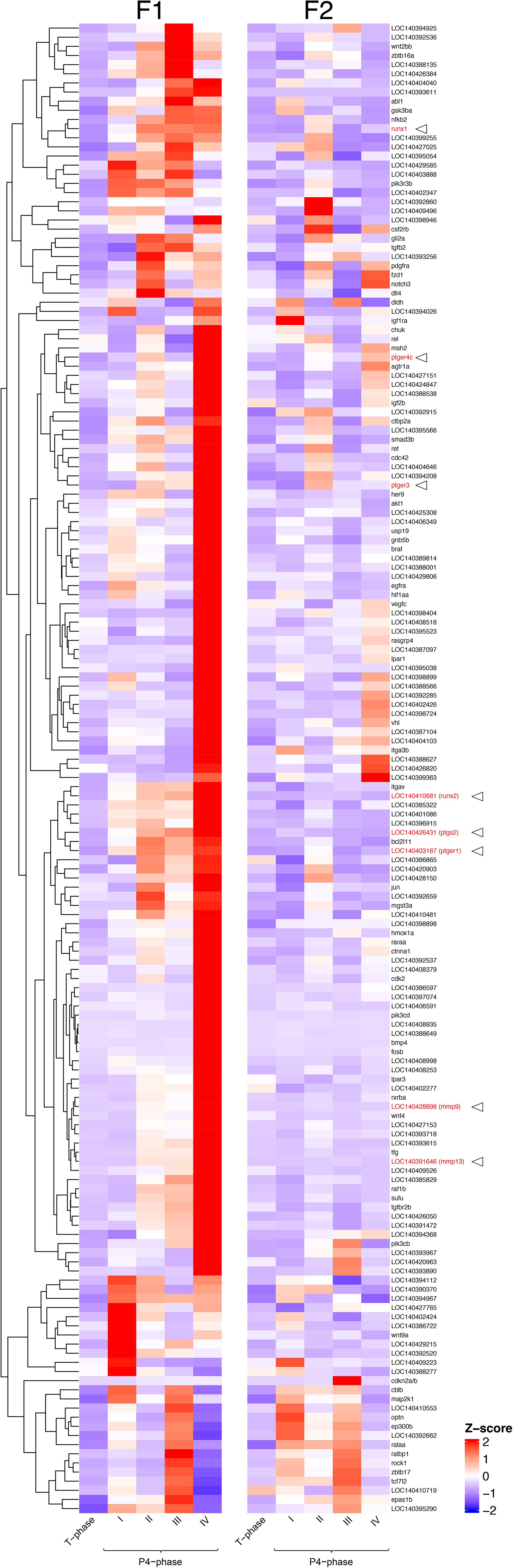
RNA-seq expression profiles of 163 candidate genes, related to Figure 2. Heatmap showing gene expression across samples with hierarchical clustering dendrograms. Samples are ordered by follicle stage (F1, left; F2, right).

**Supplementary Figure 3.**
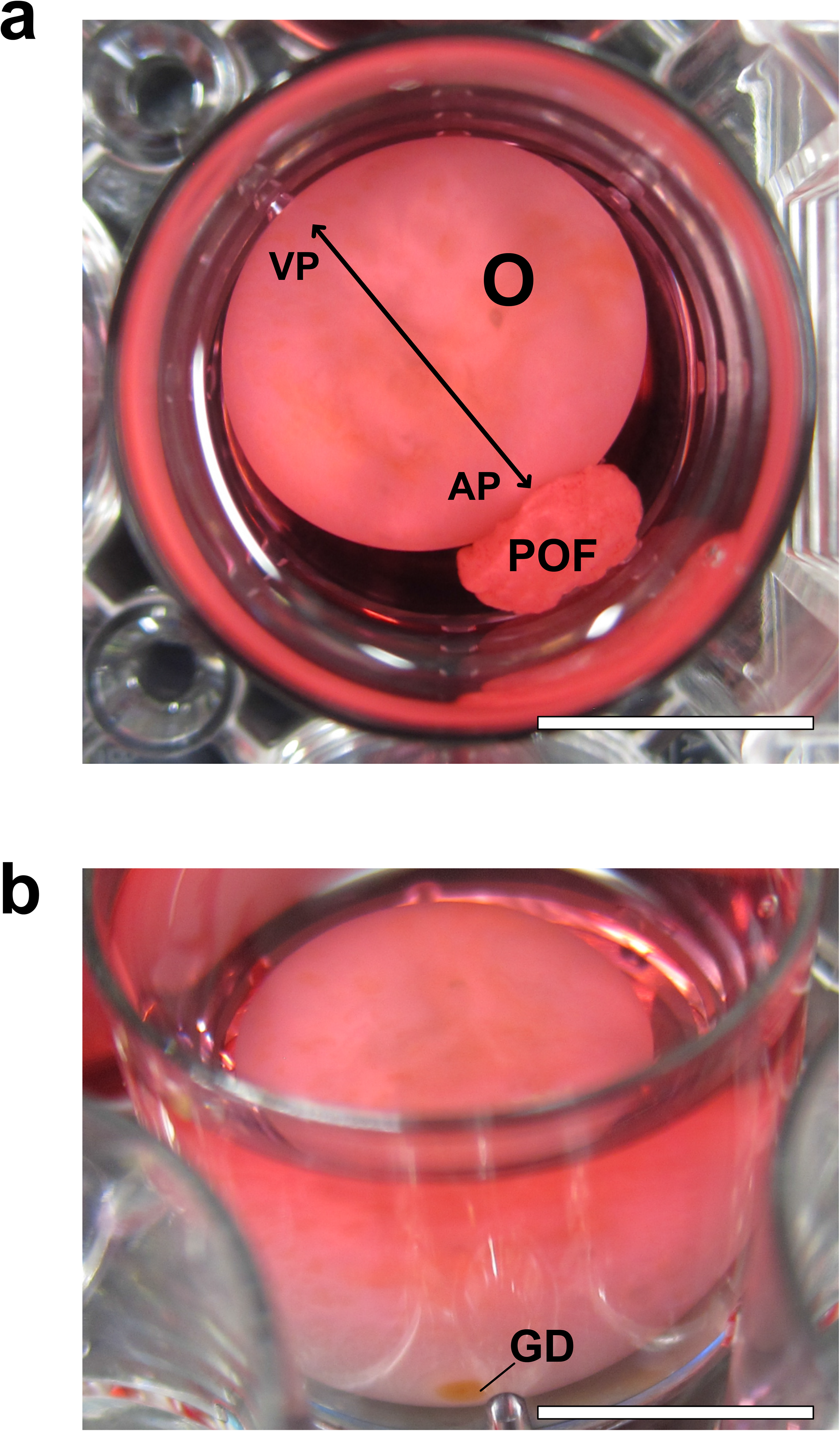
VLE-induced ovulation occurs at the vegetal pole of the follicle, related to Figure 3. a, Close-up view of a VLE-induced ovulated F1 follicle. **b,** Ovum after removal of the post-ovulatory follicular layer (POF), revealing the germinal disc (GD) located at the animal pole. VP, vegetal pole; AP, animal pole; O, ovum. Scale bar, 1 cm.

**Supplementary Figure 4.**
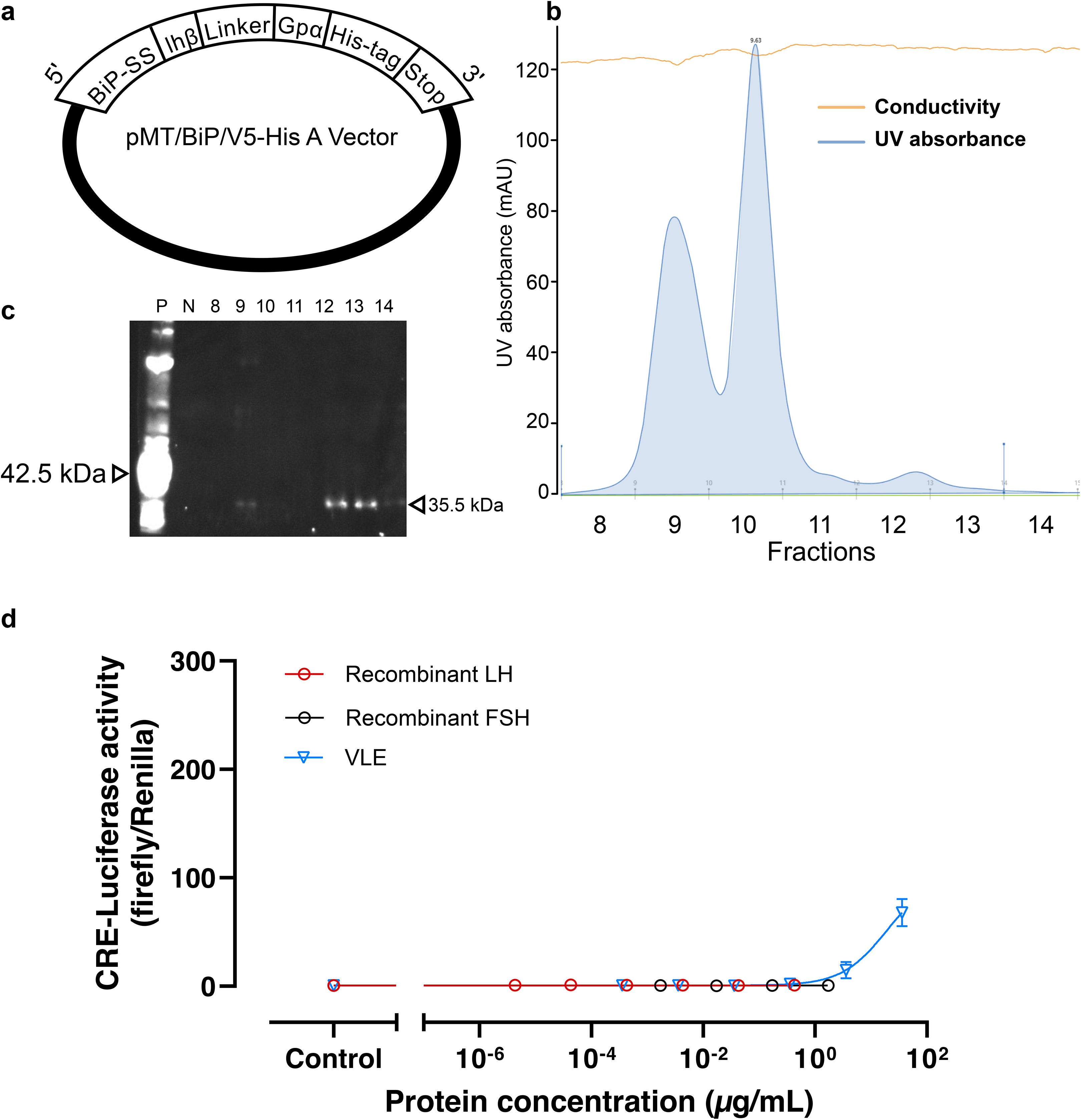
Production and purification of recombinant LH and FSH, and their activity on catshark FSH receptors, related to Figure 4. a, Expression constructs for recombinant LH (rLH) and FSH (rFSH). The LH β-subunit was replaced with FSH β-subunit to generate rFSH. b, HPLC profile of purified rLH fractions obtained after affinity and gel filtration chromatography. c, Western blot analysis of the HPLC fractions shown in b, using an anti-6×His tag antibody. d, Reporter assay using cloudy catshark FSH receptor (FSHR) expressed in HEK293A cells. Cells were treated with recombinant hormones or endogenous VLE, and receptor activation was assessed by CRE-luciferase activity.

**Supplementary Figure 5.**
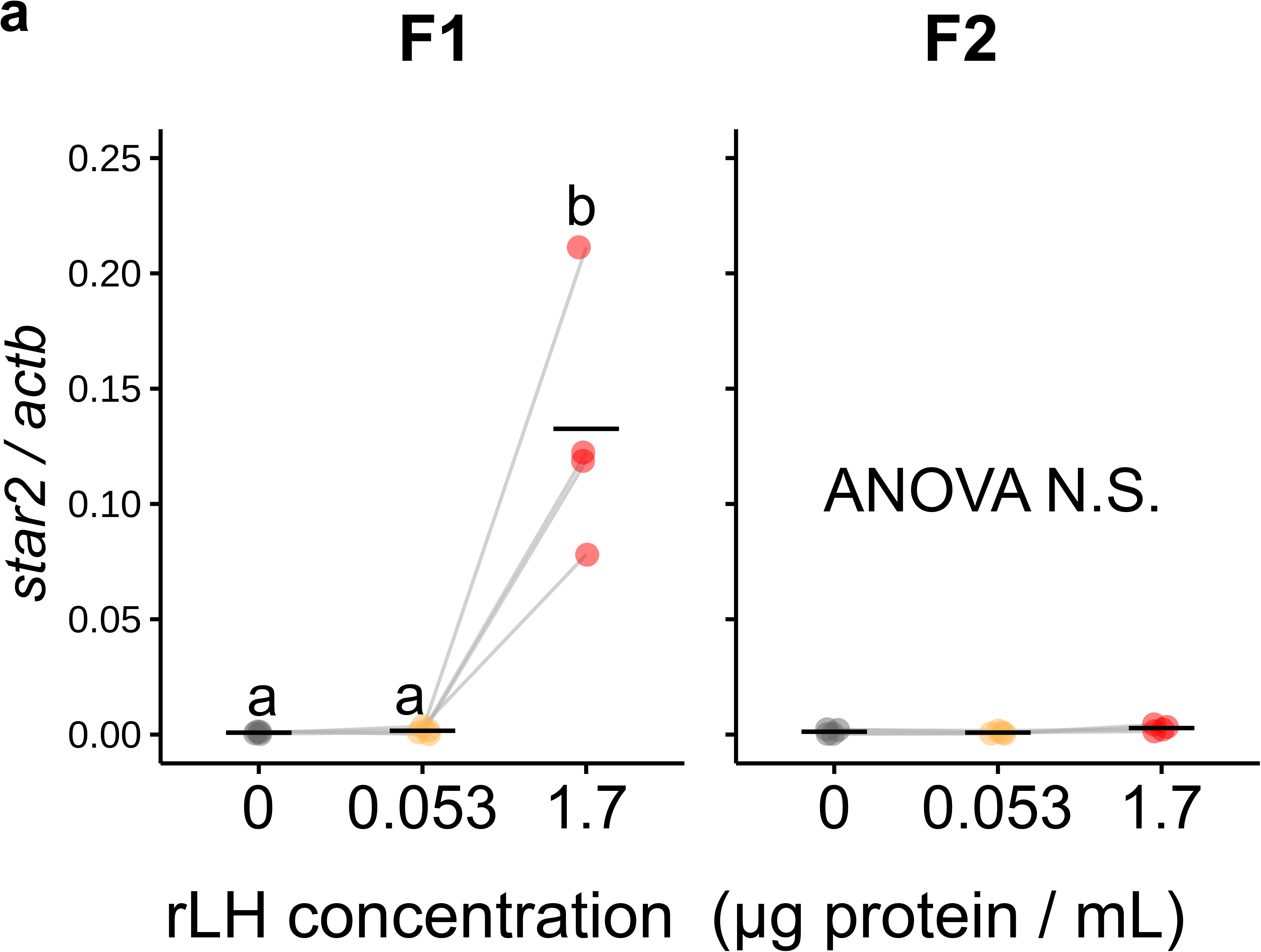
LH induces star2 expression exclusively in F1 follicles, related to Figure 4. Expression of *star2* in follicular layer strip cultures of F1 and F2 follicles after 48 h of rLH treatment, normalized to *actb*. Significant differences (*P* < 0.05) were determined by repeated-measures ANOVA followed by Tukey’s multiple comparison test and are indicated by different letters.

**Supplementary Figure 6.**
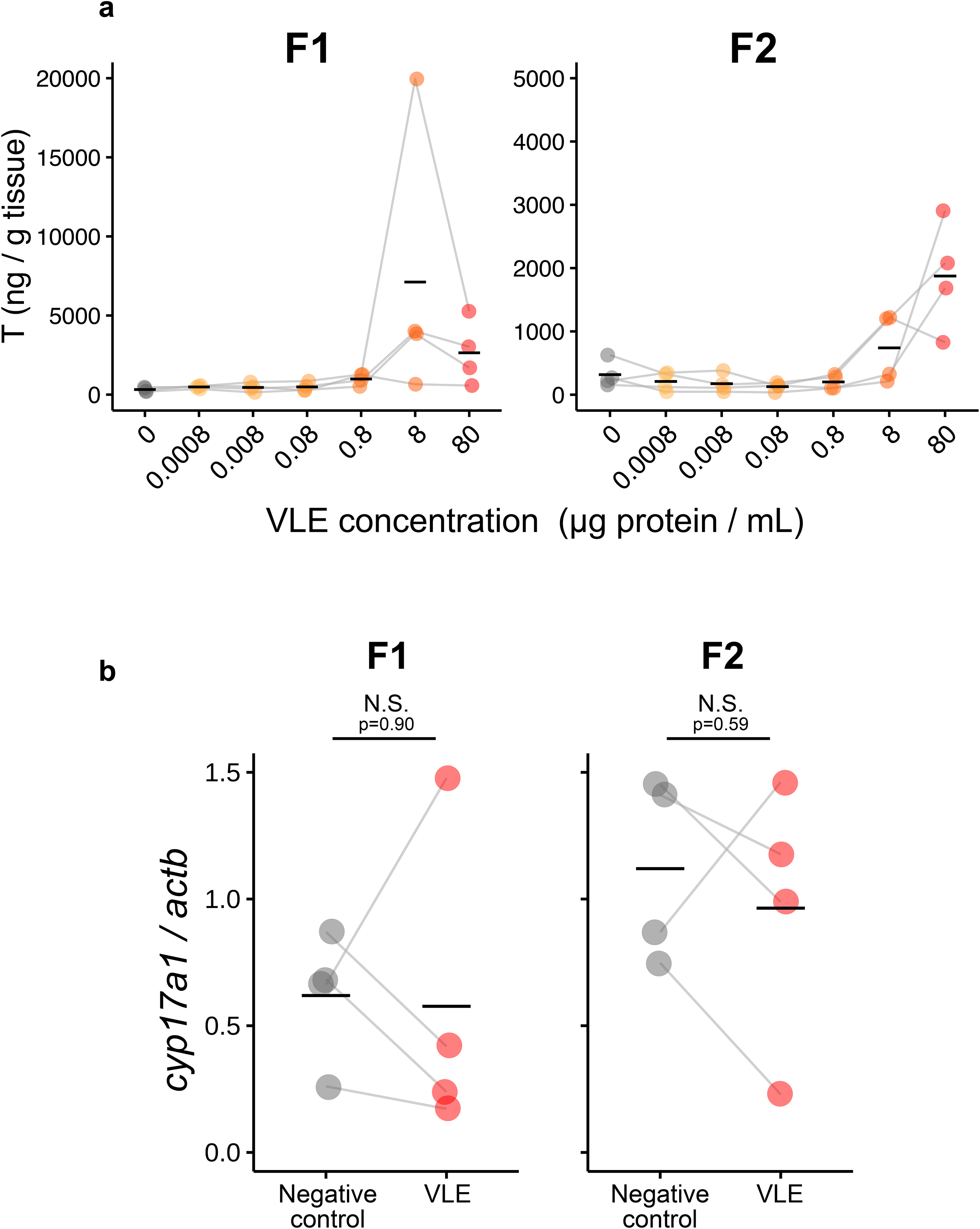
VLE treatment increases T production in both F1 and F2 follicles, related to Figure 4. a, T secretion from F1 and F2 follicular layer strips treated with increasing concentrations of VLE. Significant differences (*P* < 0.05) were determined by repeated-measures ANOVA followed by Tukey’s multiple comparison test and are indicated by different letters. b, Relative expression of *cyp17a1* in F1 and F2 follicles treated with 0 or 80 µg/mL VLE, normalized to *actb*. Statistical significance was determined by paired Student’s *t*-test. *P* < 0.05.

**Supplementary Figure 7.**
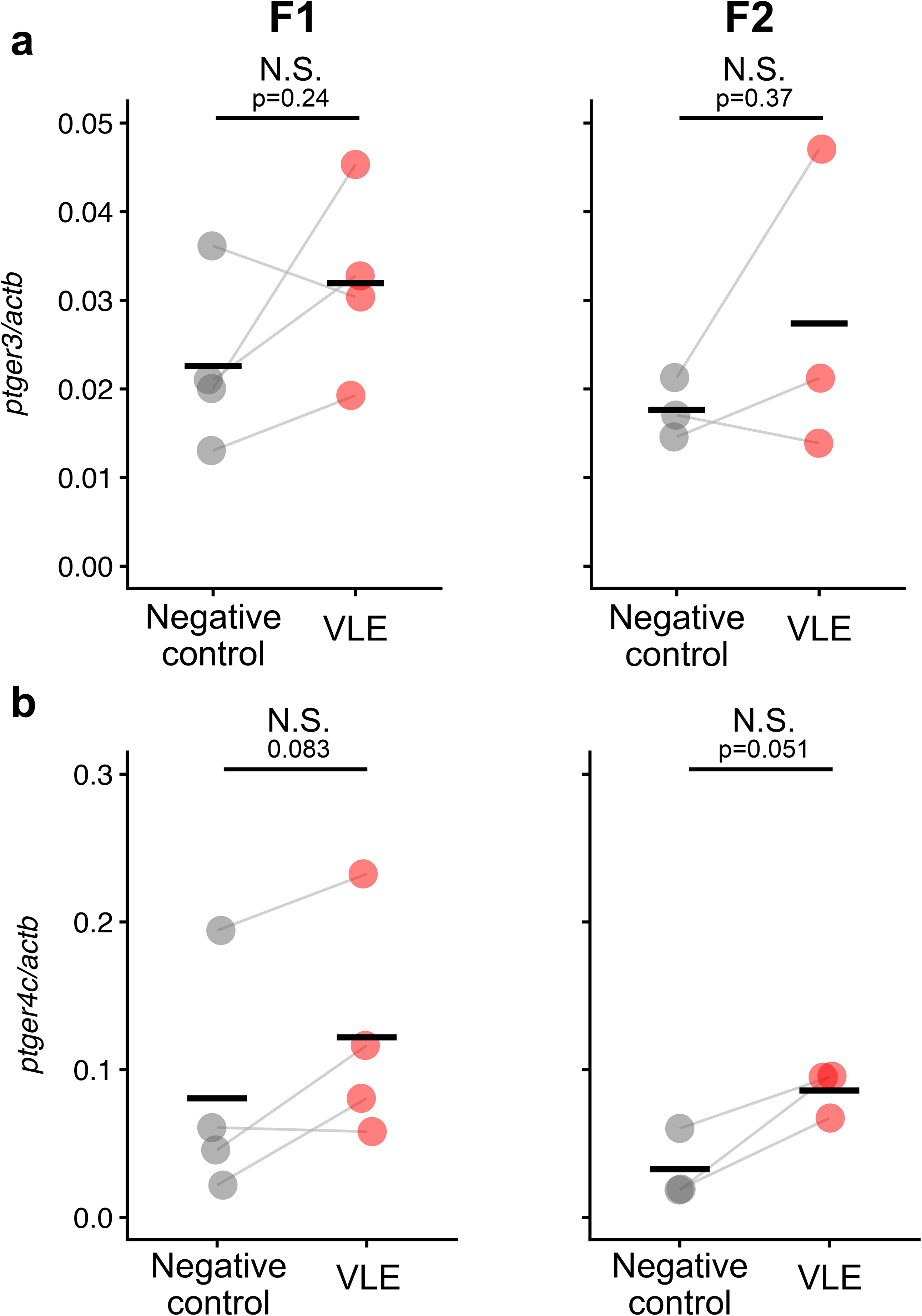
p*t*ger3 and *ptger4c* expression were not induced by VLE treatment in F1 and F2 follicles, related to Figure 5. a, b, Relative expression of *ptger3* (a) and *ptger4c* (b) in F1 and F2 follicles treated with 0 or 80 µg/ml VLE, normalized to *actb*. Statistical significance was determined by paired Student’s *t*-test. *P* < 0.05.

**Supplementary Figure 8.**
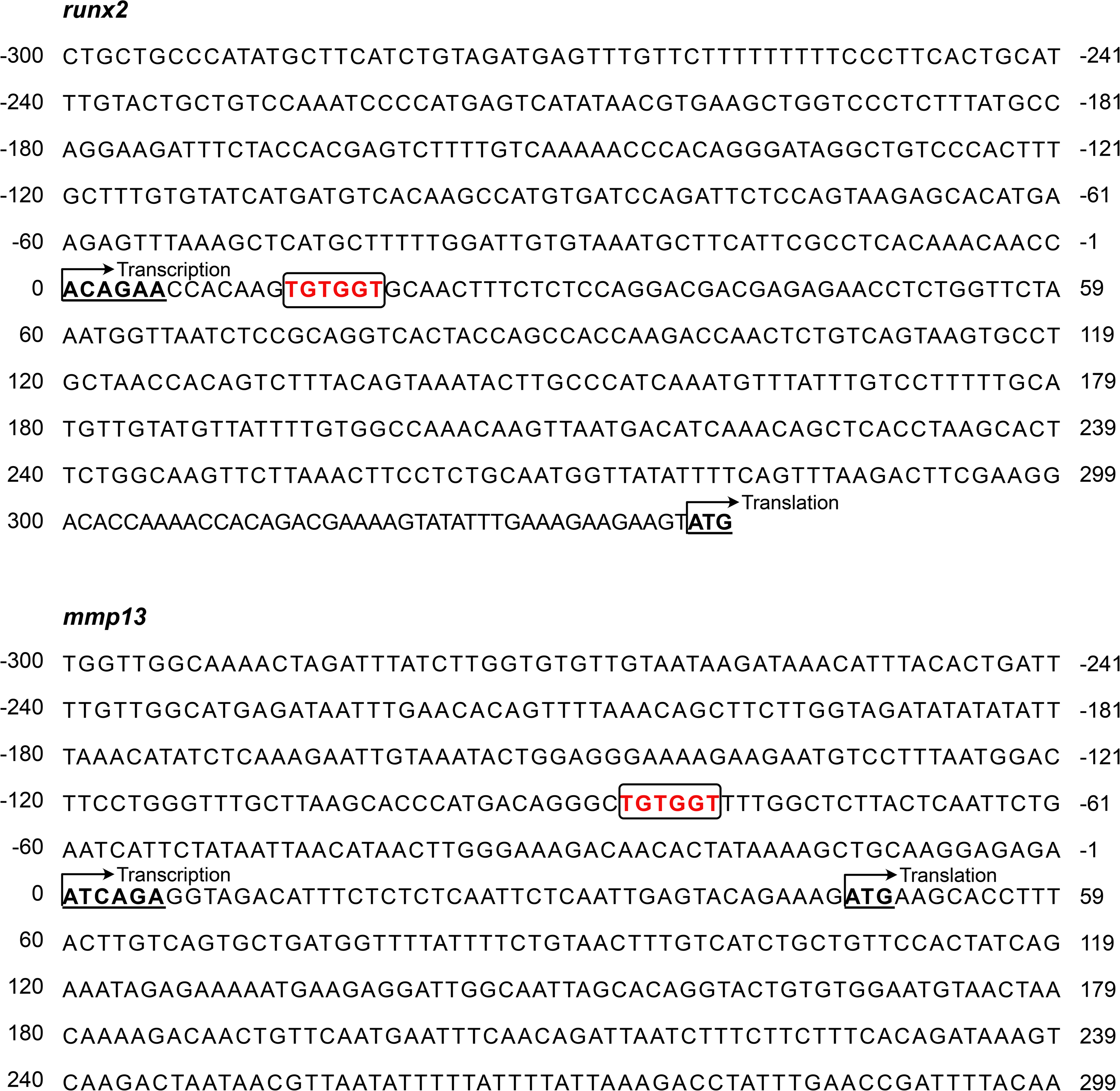
RUNX-binding motifs are present near the transcription start sites of *runx2* and *mmp13*, related to Figure 5. Schematic representation of the promoter regions of *runx2* and *mmp13*, showing the transcription start site (TSS) and putative RUNX-binding motifs. Conserved RUNX-binding sequences (5′ -TGTGGT-3′) were identified at +13 bp and −87 bp relative to the TSS of *runx2* and *mmp13*, respectively, and are highlighted in red.

**Supplementary Table 1.**
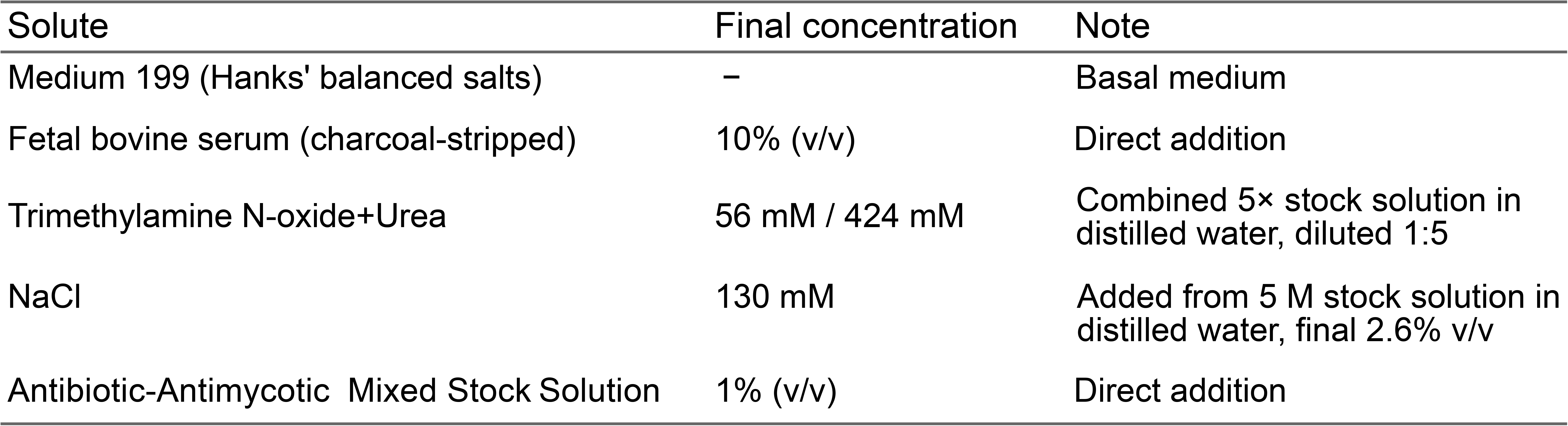
Composition of M199-based culture medium.

**Supplementary Table 2.**
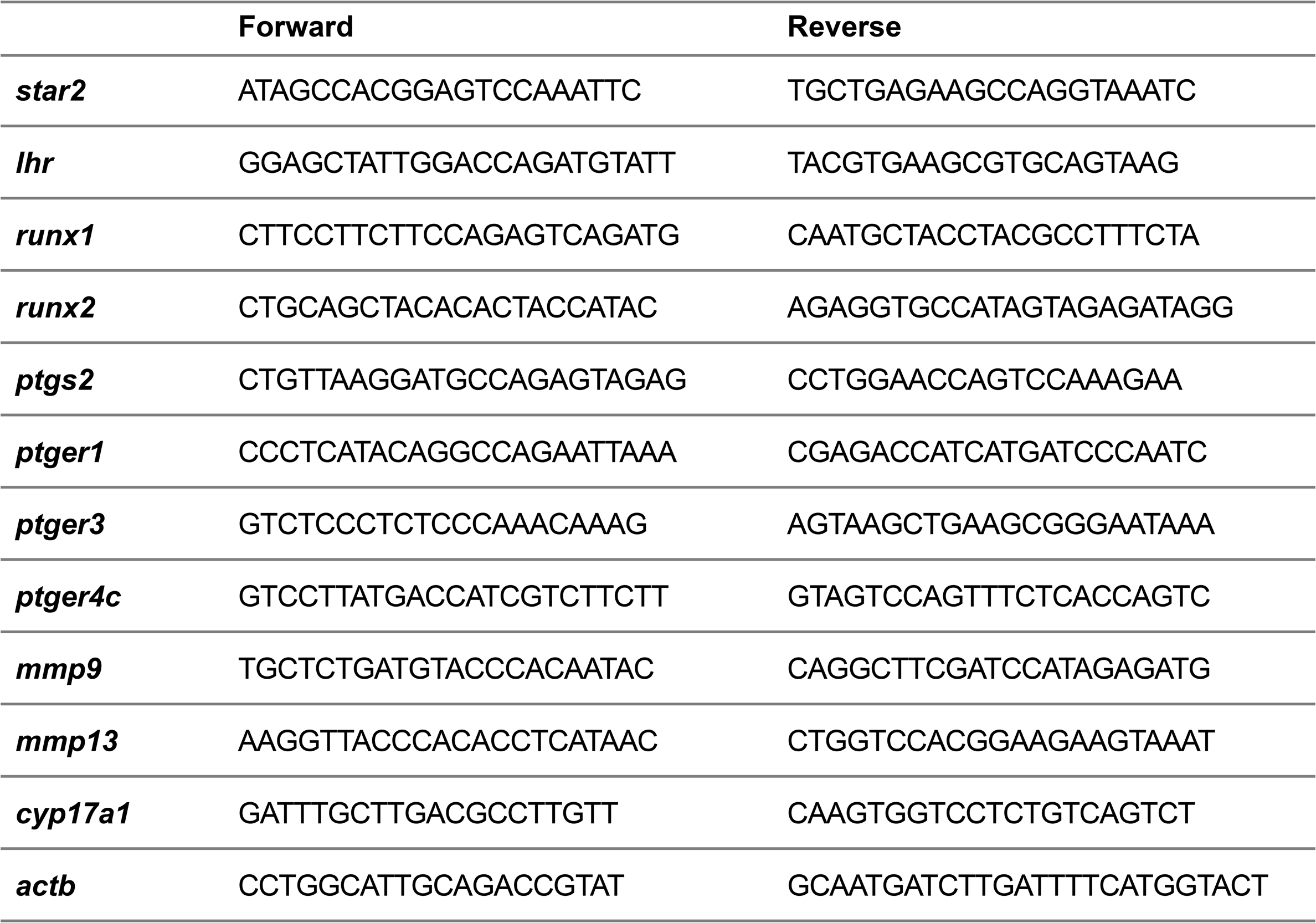
Gene specific primers used for qRT-PCR.

**Supplementary Table 3.**
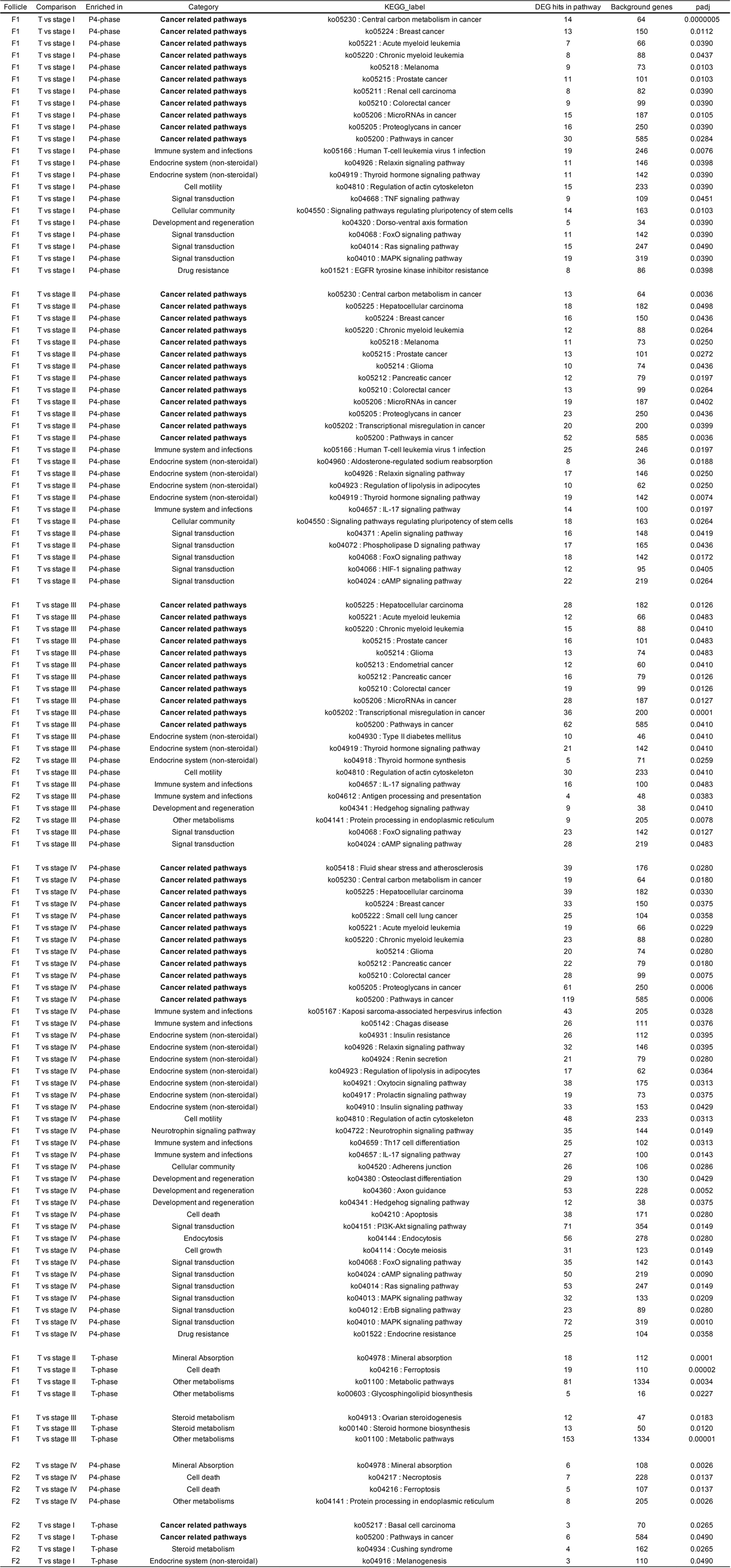
KEGG pathway enrichment results of differentially expressed genes identified from RNA-seq analysis.

