## Supplementary Figure1-8, Supplementary Table 1-3 for "cAMP responsiveness determines follicular ovulatory fate downstream of the LH receptor in the cloudy catshark"

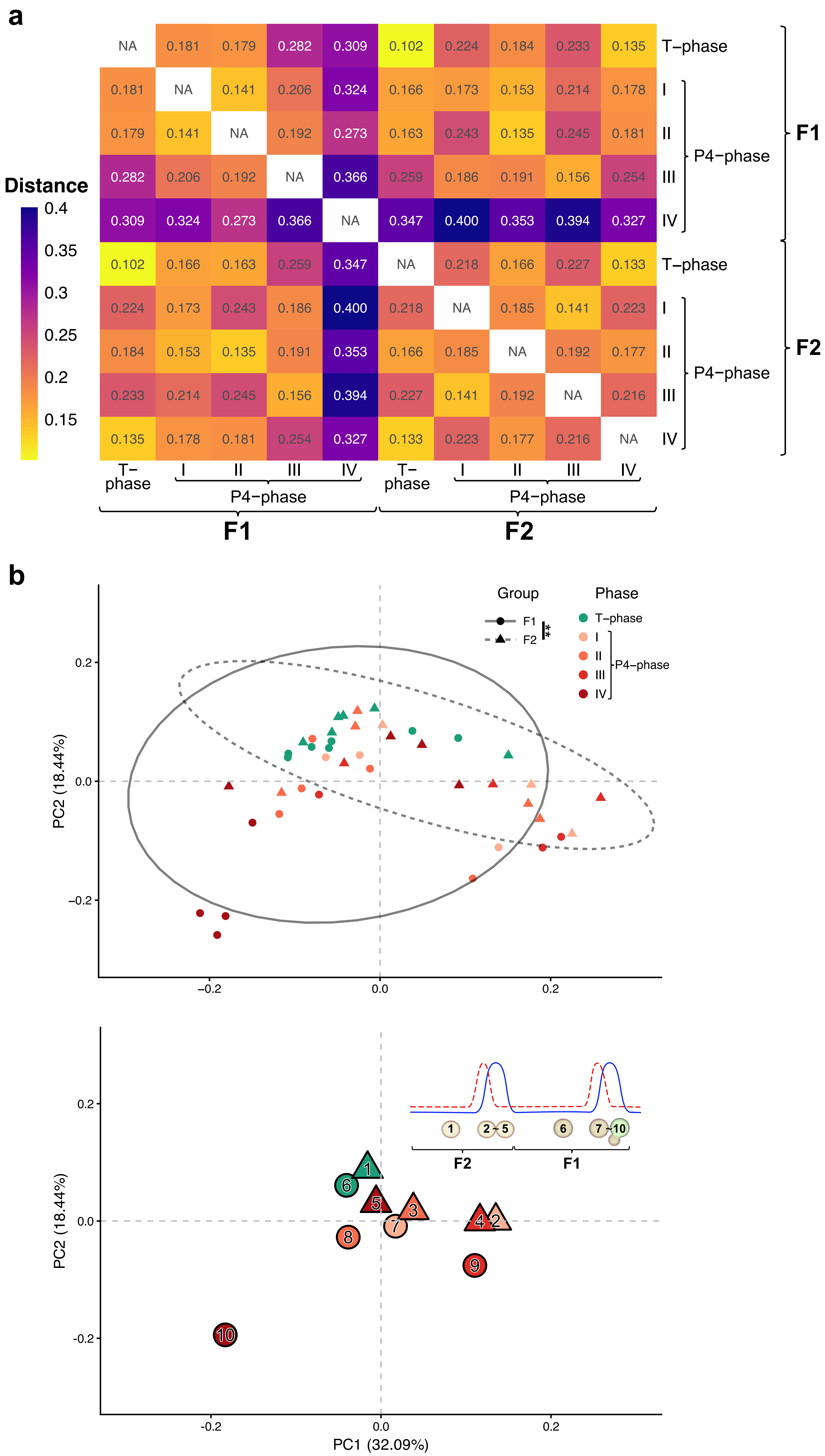

### Supplementary Figure 1

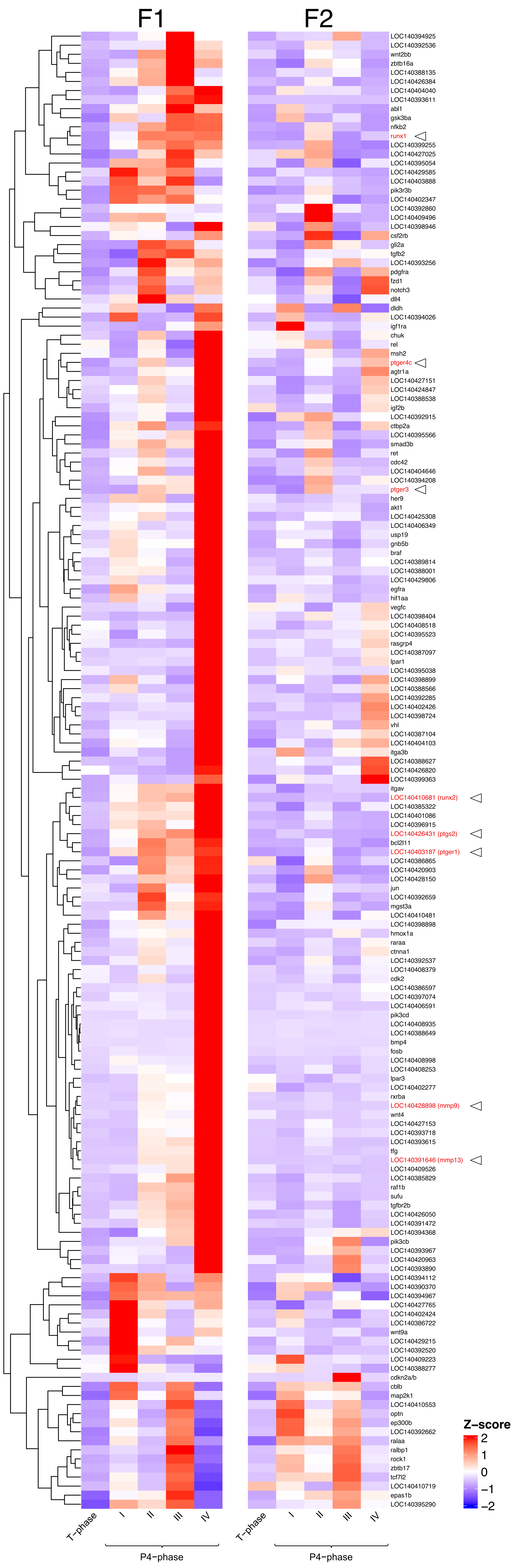

Supplementary Figure 2

**a**

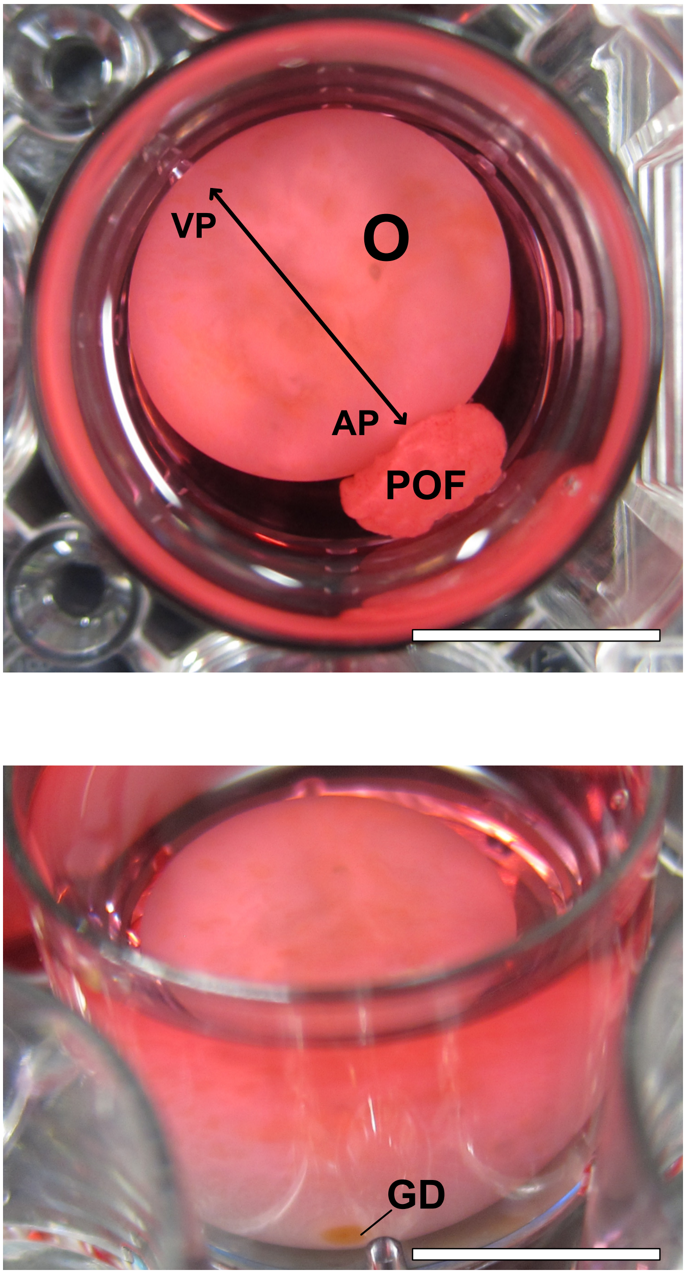

**b**

### Supplementary Figure 3

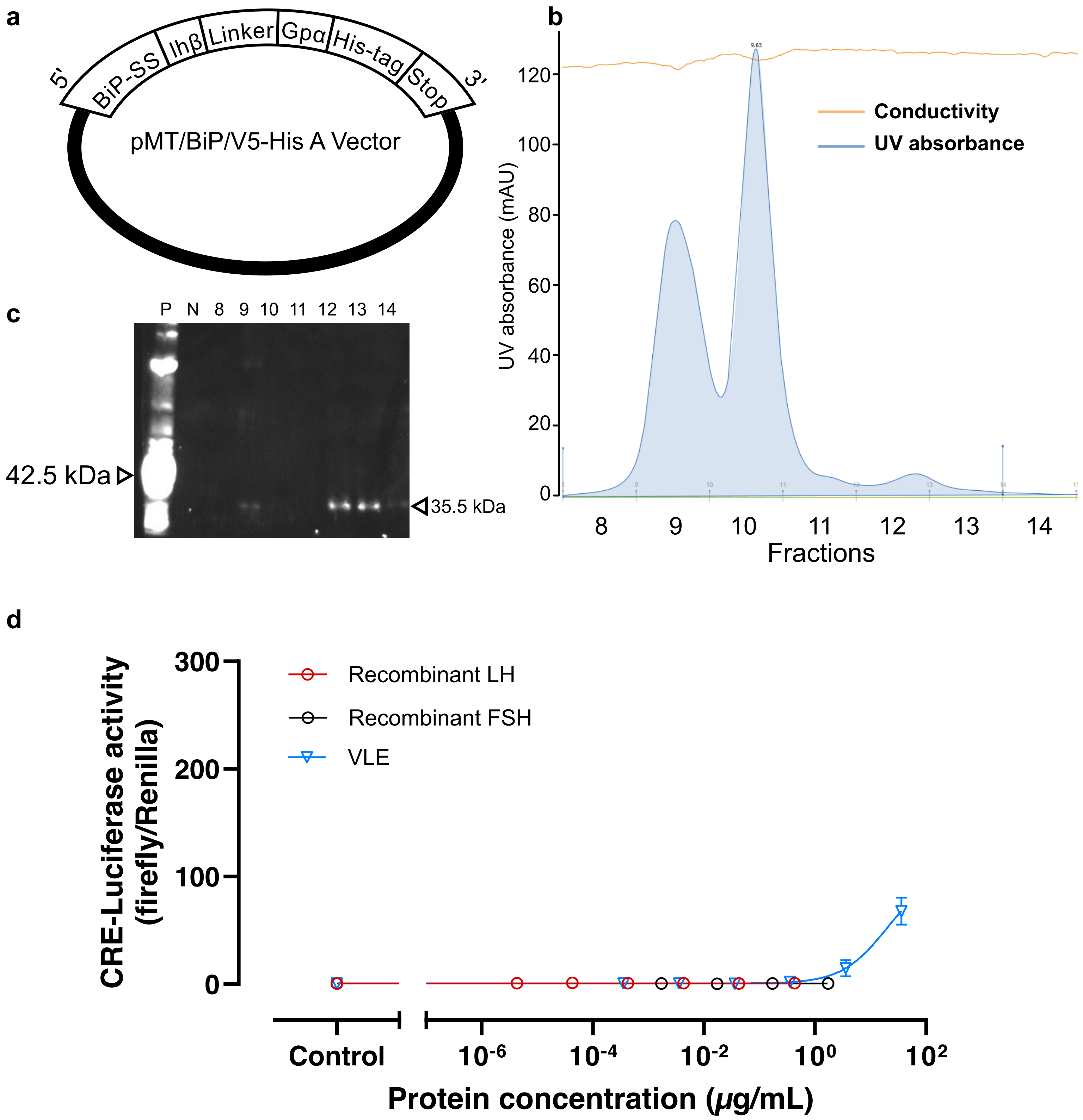

### Supplementary Figure 4

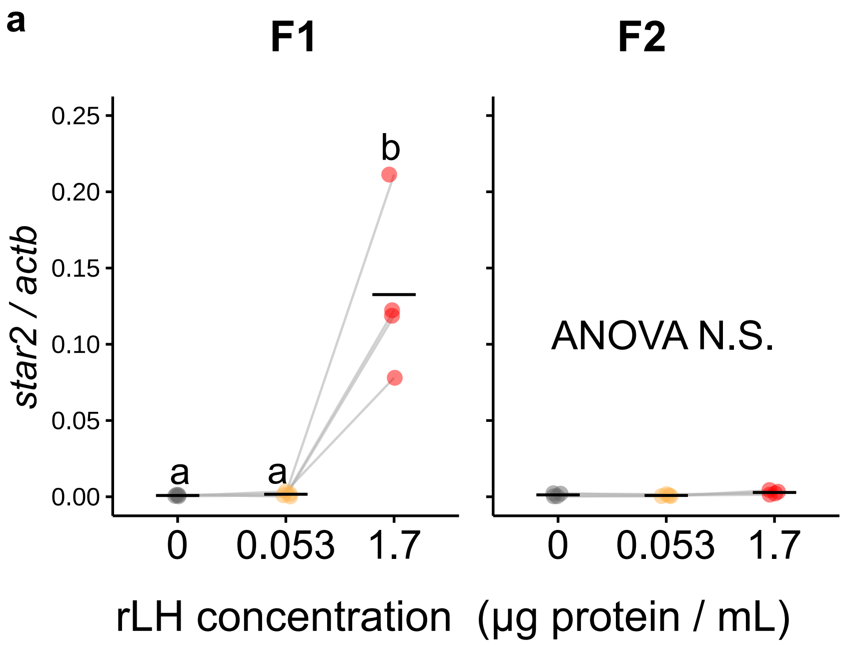

### Supplementary Figure 5

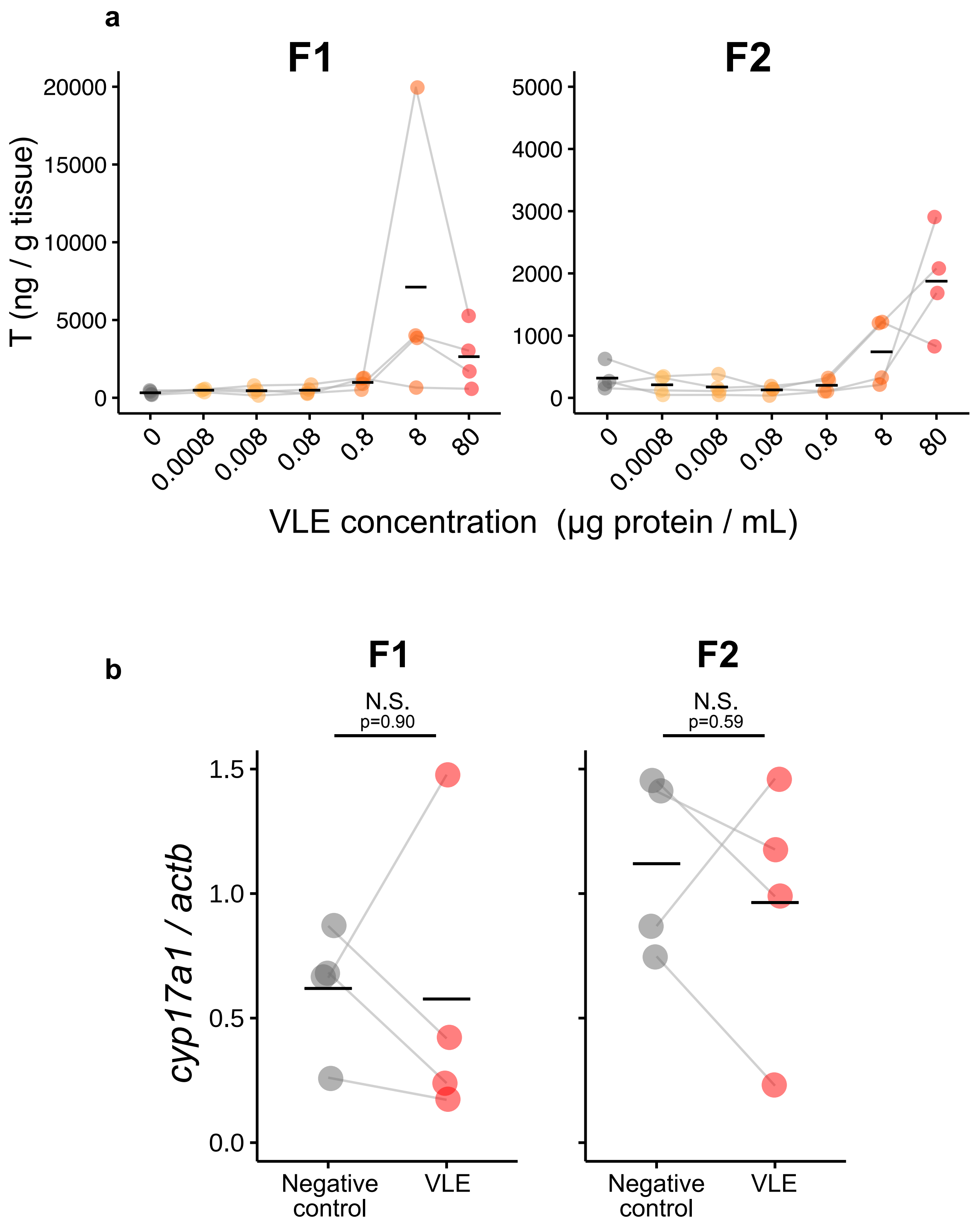

### Supplementary Figure 6

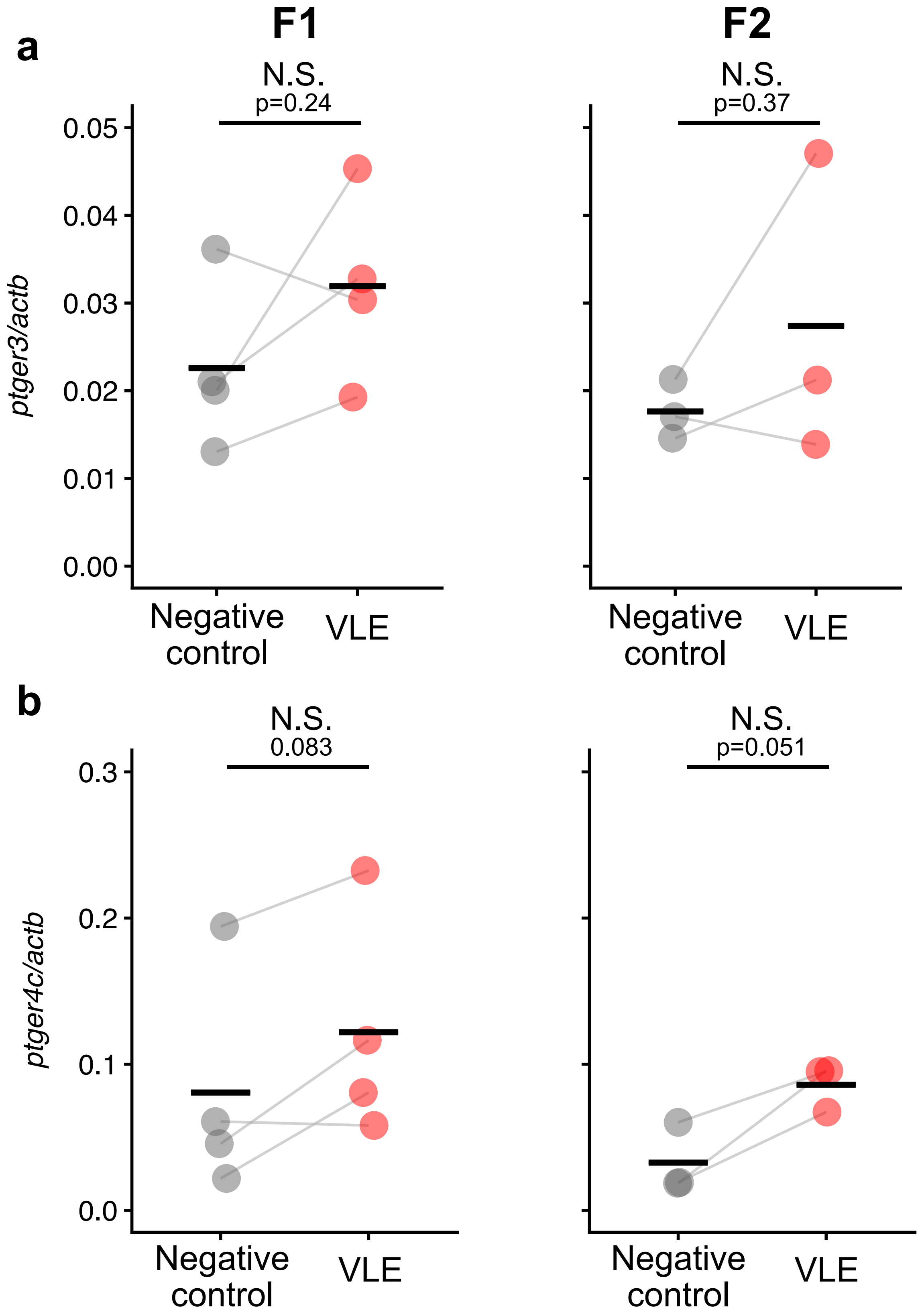

### Supplementary Figure 7

***runx2***

-300 CTGCTGCCCATATGCTTCATCTGTAGATGAGTTTGTTCTTTTTTTTTCCCTTCACTGCAT -241

-240 TTGTACTGCTGTCCAAATCCCCATGAGTCATATAACGTGAAGCTGGTCCCTCTTTATGCC -181

-180 AGGAAGATTTCTACCACGAGTCTTTTGTCAAAAACCCACAGGGATAGGCTGTCCCACTTT -121

-120 GCTTTGTGTATCATGATGTCACAAGCCATGTGATCCAGATTCTCCAGTAAGAGCACATGA -61

-60 AGAGTTTAAAGCTCATGCTTTTTGGATTGTGTAAATGCTTCATTCGCCTCACAAACAACC -1

Transcription

0 **ACAGAA**CCACAAG**TGTGGT**GCAACTTTCTCTCCAGGACGACGAGAGAACCTCTGGTTCTA 59

60 AATGGTTAATCTCCGCAGGTCACTACCAGCCACCAAGACCAACTCTGTCAGTAAGTGCCT 119

120 GCTAACCACAGTCTTTACAGTAAATACTTGCCCATCAAATGTTTATTTGTCCTTTTTGCA 179

180 TGTTGTATGTTATTTTGTGGCCAAACAAGTTAATGACATCAAACAGCTCACCTAAGCACT 239

240 TCTGGCAAGTTCTTAAACTTCCTCTGCAATGGTTATATTTTCAGTTTAAGACTTCGAAGG 299

Translation

300 ACACCAAAACCACAGACGAAAAGTATATTTGAAAGAAGAAGT**ATG**

***mmp13***

-300 TGGTTGGCAAAACTAGATTTATCTTGGTGTGTTGTAATAAGATAAACATTTACACTGATT -241

-240 TTGTTGGCATGAGATAATTTGAACACAGTTTTAAACAGCTTCTTGGTAGATATATATATT -181

-180 TAAACATATCTCAAAGAATTGTAAATACTGGAGGGAAAAGAAGAATGTCCTTTAATGGAC -121

-120 TTCCTGGGTTTGCTTAAGCACCCATGACAGGGC**TGTGGT**TTTGGCTCTTACTCAATTCTG -61

-60 AATCATTCTATAATTAACATAACTTGGGAAAGACAACACTATAAAAGCTGCAAGGAGAGA -1

Transcription Translation

0 **ATCAGA**GGTAGACATTTCTCTCTCAATTCTCAATTGAGTACAGAAAG**ATG**AAGCACCTTT 59

60 ACTTGTCAGTGCTGATGGTTTTATTTTCTGTAACTTTGTCATCTGCTGTTCCACTATCAG 119

120 AAATAGAGAAAAATGAAGAGGATTGGCAATTAGCACAGGTACTGTGTGGAATGTAACTAA 179

180 CAAAAGACAACTGTTCAATGAATTTCAACAGATTAATCTTTCTTCTTTCACAGATAAAGT 239

240 CAAGACTAATAACGTTAATATTTTTATTTTATTAAAGACCTATTTGAACCGATTTTACAA 299

### Supplementary Figure 8

**Supplementary Table 1** Composition of M199-based culture medium

| Solute | Final concentration | Note |
|---|---|---|
| Medium 199 (Hanks' balanced salts) | − | Basal medium |
| Fetal bovine serum (charcoal-stripped) | 10% (v/v) | Direct addition |
| Trimethylamine N-oxide+Urea | 56 mM / 424 mM | Combined 5× stock solution in distilled water, diluted 1:5 |
| NaCl | 130 mM | Added from 5 M stock solution in distilled water, final 2.6% v/v |
| Antibiotic-Antimycotic Mixed Stock Solution | 1% (v/v) | Direct addition |

**Supplementary Table 2** Gene specific primers used for qRT-PCR.

| | Forward | Reverse |
|---|---|---|
| ***star2*** | ATAGCCACGGAGTCCAAATTC | TGCTGAGAAGCCAGGTAAATC |
| ***lhr*** | GGAGCTATTGGACCAGATGTATT | TACGTGAAGCGTGCAGTAAG |
| ***runx1*** | CTTCCTTCTTCCAGAGTCAGATG | CAATGCTACCTACGCCTTTCTA |
| ***runx2*** | CTGCAGCTACACACTACCATAC | AGAGGTGCCATAGTAGAGATAGG |
| ***ptgs2*** | CTGTTAAGGATGCCAGAGTAGAG | CCTGGAACCAGTCCAAAGAA |
| ***ptger1*** | CCCTCATACAGGCCAGAATTAAA | CGAGACCATCATGATCCCAATC |
| ***ptger3*** | GTCTCCCTCTCCCAAACAAAG | AGTAAGCTGAAGCGGGAATAAA |
| ***ptger4c*** | GTCCTTATGACCATCGTCTTCTT | GTAGTCCAGTTTCTCACCAGTC |
| ***mmp9*** | TGCTCTGATGTACCCACAATAC | CAGGCTTCGATCCATAGAGATG |
| ***mmp13*** | AAGGTTACCCACACCTCATAAC | CTGGTCCACGGAAGAAGTAAAT |
| ***cyp17a1*** | GATTTGCTTGACGCCTTGTT | CAAGTGGTCCTCTGTCAGTCT |
| ***actb*** | CCTGGCATTGCAGACCGTAT | GCAATGATCTTGATTTTCATGGTACT |

**Supplementary Table 3** KEGG pathway enrichment results of differentially expressed genes identified from RNA-seq analysis

| Follicle | Comparison | Enriched in | Category | KEGG_label | DEG hits in pathway | Background genes | padj |
|---|---|---|---|---|---|---|---|
| F1 | T vs stage I | P4-phase | **Cancer related pathways** | ko05230 : Central carbon metabolism in cancer | 14 | 64 | 0.0000005 |
| F1 | T vs stage I | P4-phase | **Cancer related pathways** | ko05224 : Breast cancer | 13 | 150 | 0.0112 |
| F1 | T vs stage I | P4-phase | **Cancer related pathways** | ko05221 : Acute myeloid leukemia | 7 | 66 | 0.0390 |
| F1 | T vs stage I | P4-phase | **Cancer related pathways** | ko05220 : Chronic myeloid leukemia | 8 | 88 | 0.0437 |
| F1 | T vs stage I | P4-phase | **Cancer related pathways** | ko05218 : Melanoma | 9 | 73 | 0.0103 |
| F1 | T vs stage I | P4-phase | **Cancer related pathways** | ko05215 : Prostate cancer | 11 | 101 | 0.0103 |
| F1 | T vs stage I | P4-phase | **Cancer related pathways** | ko05211 : Renal cell carcinoma | 8 | 82 | 0.0390 |
| F1 | T vs stage I | P4-phase | **Cancer related pathways** | ko05210 : Colorectal cancer | 9 | 99 | 0.0390 |
| F1 | T vs stage I | P4-phase | **Cancer related pathways** | ko05206 : MicroRNAs in cancer | 15 | 187 | 0.0105 |
| F1 | T vs stage I | P4-phase | **Cancer related pathways** | ko05205 : Proteoglycans in cancer | 16 | 250 | 0.0390 |
| F1 | T vs stage I | P4-phase | **Cancer related pathways** | ko05200 : Pathways in cancer | 30 | 585 | 0.0284 |
| F1 | T vs stage I | P4-phase | Immune system and infections | ko05166 : Human T-cell leukemia virus 1 infection | 19 | 246 | 0.0076 |
| F1 | T vs stage I | P4-phase | Endocrine system (non-steroidal) | ko04926 : Relaxin signaling pathway | 11 | 146 | 0.0398 |
| F1 | T vs stage I | P4-phase | Endocrine system (non-steroidal) | ko04919 : Thyroid hormone signaling pathway | 11 | 142 | 0.0390 |
| F1 | T vs stage I | P4-phase | Cell motility | ko04810 : Regulation of actin cytoskeleton | 15 | 233 | 0.0390 |
| F1 | T vs stage I | P4-phase | Signal transduction | ko04668 : TNF signaling pathway | 9 | 109 | 0.0451 |
| F1 | T vs stage I | P4-phase | Cellular community | ko04550 : Signaling pathways regulating pluripotency of stem cells | 14 | 163 | 0.0103 |
| F1 | T vs stage I | P4-phase | Development and regeneration | ko04320 : Dorso-ventral axis formation | 5 | 34 | 0.0390 |
| F1 | T vs stage I | P4-phase | Signal transduction | ko04068 : FoxO signaling pathway | 11 | 142 | 0.0390 |
| F1 | T vs stage I | P4-phase | Signal transduction | ko04014 : Ras signaling pathway | 15 | 247 | 0.0490 |
| F1 | T vs stage I | P4-phase | Signal transduction | ko04010 : MAPK signaling pathway | 19 | 319 | 0.0390 |
| F1 | T vs stage I | P4-phase | Drug resistance | ko01521 : EGFR tyrosine kinase inhibitor resistance | 8 | 86 | 0.0398 |
| F1 | T vs stage II | P4-phase | **Cancer related pathways** | ko05230 : Central carbon metabolism in cancer | 13 | 64 | 0.0036 |
| F1 | T vs stage II | P4-phase | **Cancer related pathways** | ko05225 : Hepatocellular carcinoma | 18 | 182 | 0.0498 |
| F1 | T vs stage II | P4-phase | **Cancer related pathways** | ko05224 : Breast cancer | 16 | 150 | 0.0436 |
| F1 | T vs stage II | P4-phase | **Cancer related pathways** | ko05220 : Chronic myeloid leukemia | 12 | 88 | 0.0264 |
| F1 | T vs stage II | P4-phase | **Cancer related pathways** | ko05218 : Melanoma | 11 | 73 | 0.0250 |
| F1 | T vs stage II | P4-phase | **Cancer related pathways** | ko05215 : Prostate cancer | 13 | 101 | 0.0272 |
| F1 | T vs stage II | P4-phase | **Cancer related pathways** | ko05214 : Glioma | 10 | 74 | 0.0436 |
| F1 | T vs stage II | P4-phase | **Cancer related pathways** | ko05212 : Pancreatic cancer | 12 | 79 | 0.0197 |
| F1 | T vs stage II | P4-phase | **Cancer related pathways** | ko05210 : Colorectal cancer | 13 | 99 | 0.0264 |
| F1 | T vs stage II | P4-phase | **Cancer related pathways** | ko05206 : MicroRNAs in cancer | 19 | 187 | 0.0402 |
| F1 | T vs stage II | P4-phase | **Cancer related pathways** | ko05205 : Proteoglycans in cancer | 23 | 250 | 0.0436 |
| F1 | T vs stage II | P4-phase | **Cancer related pathways** | ko05202 : Transcriptional misregulation in cancer | 20 | 200 | 0.0399 |
| F1 | T vs stage II | P4-phase | **Cancer related pathways** | ko05200 : Pathways in cancer | 52 | 585 | 0.0036 |
| F1 | T vs stage II | P4-phase | Immune system and infections | ko05166 : Human T-cell leukemia virus 1 infection | 25 | 246 | 0.0197 |
| F1 | T vs stage II | P4-phase | Endocrine system (non-steroidal) | ko04960 : Aldosterone-regulated sodium reabsorption | 8 | 36 | 0.0188 |
| F1 | T vs stage II | P4-phase | Endocrine system (non-steroidal) | ko04926 : Relaxin signaling pathway | 17 | 146 | 0.0250 |
| F1 | T vs stage II | P4-phase | Endocrine system (non-steroidal) | ko04923 : Regulation of lipolysis in adipocytes | 10 | 62 | 0.0250 |
| F1 | T vs stage II | P4-phase | Endocrine system (non-steroidal) | ko04919 : Thyroid hormone signaling pathway | 19 | 142 | 0.0074 |
| F1 | T vs stage II | P4-phase | Immune system and infections | ko04657 : IL-17 signaling pathway | 14 | 100 | 0.0197 |
| F1 | T vs stage II | P4-phase | Cellular community | ko04550 : Signaling pathways regulating pluripotency of stem cells | 18 | 163 | 0.0264 |
| F1 | T vs stage II | P4-phase | Signal transduction | ko04371 : Apelin signaling pathway | 16 | 148 | 0.0419 |
| F1 | T vs stage II | P4-phase | Signal transduction | ko04072 : Phospholipase D signaling pathway | 17 | 165 | 0.0436 |
| F1 | T vs stage II | P4-phase | Signal transduction | ko04068 : FoxO signaling pathway | 18 | 142 | 0.0172 |
| F1 | T vs stage II | P4-phase | Signal transduction | ko04066 : HIF-1 signaling pathway | 12 | 95 | 0.0405 |
| F1 | T vs stage II | P4-phase | Signal transduction | ko04024 : cAMP signaling pathway | 22 | 219 | 0.0264 |
| F1 | T vs stage III | P4-phase | **Cancer related pathways** | ko05225 : Hepatocellular carcinoma | 28 | 182 | 0.0126 |
| F1 | T vs stage III | P4-phase | **Cancer related pathways** | ko05221 : Acute myeloid leukemia | 12 | 66 | 0.0483 |
| F1 | T vs stage III | P4-phase | **Cancer related pathways** | ko05220 : Chronic myeloid leukemia | 15 | 88 | 0.0410 |
| F1 | T vs stage III | P4-phase | **Cancer related pathways** | ko05215 : Prostate cancer | 16 | 101 | 0.0483 |
| F1 | T vs stage III | P4-phase | **Cancer related pathways** | ko05214 : Glioma | 13 | 74 | 0.0483 |
| F1 | T vs stage III | P4-phase | **Cancer related pathways** | ko05213 : Endometrial cancer | 12 | 60 | 0.0410 |
| F1 | T vs stage III | P4-phase | **Cancer related pathways** | ko05212 : Pancreatic cancer | 16 | 79 | 0.0126 |
| F1 | T vs stage III | P4-phase | **Cancer related pathways** | ko05210 : Colorectal cancer | 19 | 99 | 0.0126 |
| F1 | T vs stage III | P4-phase | **Cancer related pathways** | ko05206 : MicroRNAs in cancer | 28 | 187 | 0.0127 |
| F1 | T vs stage III | P4-phase | **Cancer related pathways** | ko05202 : Transcriptional misregulation in cancer | 36 | 200 | 0.0001 |
| F1 | T vs stage III | P4-phase | **Cancer related pathways** | ko05200 : Pathways in cancer | 62 | 585 | 0.0410 |
| F1 | T vs stage III | P4-phase | Endocrine system (non-steroidal) | ko04930 : Type II diabetes mellitus | 10 | 46 | 0.0410 |
| F1 | T vs stage III | P4-phase | Endocrine system (non-steroidal) | ko04919 : Thyroid hormone signaling pathway | 21 | 142 | 0.0410 |
| F2 | T vs stage III | P4-phase | Endocrine system (non-steroidal) | ko04918 : Thyroid hormone synthesis | 5 | 71 | 0.0259 |
| F1 | T vs stage III | P4-phase | Cell motility | ko04810 : Regulation of actin cytoskeleton | 30 | 233 | 0.0410 |
| F1 | T vs stage III | P4-phase | Immune system and infections | ko04657 : IL-17 signaling pathway | 16 | 100 | 0.0483 |
| F2 | T vs stage III | P4-phase | Immune system and infections | ko04612 : Antigen processing and presentation | 4 | 48 | 0.0383 |
| F1 | T vs stage III | P4-phase | Development and regeneration | ko04341 : Hedgehog signaling pathway | 9 | 38 | 0.0410 |
| F2 | T vs stage III | P4-phase | Other metabolisms | ko04141 : Protein processing in endoplasmic reticulum | 9 | 205 | 0.0078 |
| F1 | T vs stage III | P4-phase | Signal transduction | ko04068 : FoxO signaling pathway | 23 | 142 | 0.0127 |
| F1 | T vs stage III | P4-phase | Signal transduction | ko04024 : cAMP signaling pathway | 28 | 219 | 0.0483 |
| F1 | T vs stage IV | P4-phase | **Cancer related pathways** | ko05418 : Fluid shear stress and atherosclerosis | 39 | 176 | 0.0280 |
| F1 | T vs stage IV | P4-phase | **Cancer related pathways** | ko05230 : Central carbon metabolism in cancer | 19 | 64 | 0.0180 |
| F1 | T vs stage IV | P4-phase | **Cancer related pathways** | ko05225 : Hepatocellular carcinoma | 39 | 182 | 0.0330 |
| F1 | T vs stage IV | P4-phase | **Cancer related pathways** | ko05224 : Breast cancer | 33 | 150 | 0.0375 |
| F1 | T vs stage IV | P4-phase | **Cancer related pathways** | ko05222 : Small cell lung cancer | 25 | 104 | 0.0358 |
| F1 | T vs stage IV | P4-phase | **Cancer related pathways** | ko05221 : Acute myeloid leukemia | 19 | 66 | 0.0229 |
| F1 | T vs stage IV | P4-phase | **Cancer related pathways** | ko05220 : Chronic myeloid leukemia | 23 | 88 | 0.0280 |
| F1 | T vs stage IV | P4-phase | **Cancer related pathways** | ko05214 : Glioma | 20 | 74 | 0.0280 |
| F1 | T vs stage IV | P4-phase | **Cancer related pathways** | ko05212 : Pancreatic cancer | 22 | 79 | 0.0180 |
| F1 | T vs stage IV | P4-phase | **Cancer related pathways** | ko05210 : Colorectal cancer | 28 | 99 | 0.0075 |
| F1 | T vs stage IV | P4-phase | **Cancer related pathways** | ko05205 : Proteoglycans in cancer | 61 | 250 | 0.0006 |
| F1 | T vs stage IV | P4-phase | **Cancer related pathways** | ko05200 : Pathways in cancer | 119 | 585 | 0.0006 |
| F1 | T vs stage IV | P4-phase | Immune system and infections | ko05167 : Kaposi sarcoma-associated herpesvirus infection | 43 | 205 | 0.0328 |
| F1 | T vs stage IV | P4-phase | Immune system and infections | ko05142 : Chagas disease | 26 | 111 | 0.0376 |
| F1 | T vs stage IV | P4-phase | Endocrine system (non-steroidal) | ko04931 : Insulin resistance | 26 | 112 | 0.0395 |
| F1 | T vs stage IV | P4-phase | Endocrine system (non-steroidal) | ko04926 : Relaxin signaling pathway | 32 | 146 | 0.0395 |
| F1 | T vs stage IV | P4-phase | Endocrine system (non-steroidal) | ko04924 : Renin secretion | 21 | 79 | 0.0280 |
| F1 | T vs stage IV | P4-phase | Endocrine system (non-steroidal) | ko04923 : Regulation of lipolysis in adipocytes | 17 | 62 | 0.0364 |
| F1 | T vs stage IV | P4-phase | Endocrine system (non-steroidal) | ko04921 : Oxytocin signaling pathway | 38 | 175 | 0.0313 |
| F1 | T vs stage IV | P4-phase | Endocrine system (non-steroidal) | ko04917 : Prolactin signaling pathway | 19 | 73 | 0.0375 |
| F1 | T vs stage IV | P4-phase | Endocrine system (non-steroidal) | ko04910 : Insulin signaling pathway | 33 | 153 | 0.0429 |
| F1 | T vs stage IV | P4-phase | Cell motility | ko04810 : Regulation of actin cytoskeleton | 48 | 233 | 0.0313 |
| F1 | T vs stage IV | P4-phase | Neurotrophin signaling pathway | ko04722 : Neurotrophin signaling pathway | 35 | 144 | 0.0149 |
| F1 | T vs stage IV | P4-phase | Immune system and infections | ko04659 : Th17 cell differentiation | 25 | 102 | 0.0313 |
| F1 | T vs stage IV | P4-phase | Immune system and infections | ko04657 : IL-17 signaling pathway | 27 | 100 | 0.0143 |
| F1 | T vs stage IV | P4-phase | Cellular community | ko04520 : Adherens junction | 26 | 106 | 0.0286 |
| F1 | T vs stage IV | P4-phase | Development and regeneration | ko04380 : Osteoclast differentiation | 29 | 130 | 0.0429 |
| F1 | T vs stage IV | P4-phase | Development and regeneration | ko04360 : Axon guidance | 53 | 228 | 0.0052 |
| F1 | T vs stage IV | P4-phase | Development and regeneration | ko04341 : Hedgehog signaling pathway | 12 | 38 | 0.0375 |
| F1 | T vs stage IV | P4-phase | Cell death | ko04210 : Apoptosis | 38 | 171 | 0.0280 |
| F1 | T vs stage IV | P4-phase | Signal transduction | ko04151 : PI3K-Akt signaling pathway | 71 | 354 | 0.0149 |
| F1 | T vs stage IV | P4-phase | Endocytosis | ko04144 : Endocytosis | 56 | 278 | 0.0280 |
| F1 | T vs stage IV | P4-phase | Cell growth | ko04114 : Oocyte meiosis | 31 | 123 | 0.0149 |
| F1 | T vs stage IV | P4-phase | Signal transduction | ko04068 : FoxO signaling pathway | 35 | 142 | 0.0143 |
| F1 | T vs stage IV | P4-phase | Signal transduction | ko04024 : cAMP signaling pathway | 50 | 219 | 0.0090 |
| F1 | T vs stage IV | P4-phase | Signal transduction | ko04014 : Ras signaling pathway | 53 | 247 | 0.0149 |
| F1 | T vs stage IV | P4-phase | Signal transduction | ko04013 : MAPK signaling pathway | 32 | 133 | 0.0209 |
| F1 | T vs stage IV | P4-phase | Signal transduction | ko04012 : ErbB signaling pathway | 23 | 89 | 0.0280 |
| F1 | T vs stage IV | P4-phase | Signal transduction | ko04010 : MAPK signaling pathway | 72 | 319 | 0.0010 |
| F1 | T vs stage IV | P4-phase | Drug resistance | ko01522 : Endocrine resistance | 25 | 104 | 0.0358 |
| F1 | T vs stage II | T-phase | Mineral Absorption | ko04978 : Mineral absorption | 18 | 112 | 0.0001 |
| F1 | T vs stage II | T-phase | Cell death | ko04216 : Ferroptosis | 19 | 110 | 0.00002 |
| F1 | T vs stage II | T-phase | Other metabolisms | ko01100 : Metabolic pathways | 81 | 1334 | 0.0034 |
| F1 | T vs stage II | T-phase | Other metabolisms | ko00603 : Glycosphingolipid biosynthesis | 5 | 16 | 0.0227 |
| F1 | T vs stage III | T-phase | Steroid metabolism | ko04913 : Ovarian steroidogenesis | 12 | 47 | 0.0183 |
| F1 | T vs stage III | T-phase | Steroid metabolism | ko00140 : Steroid hormone biosynthesis | 13 | 50 | 0.0120 |
| F1 | T vs stage III | T-phase | Other metabolisms | ko01100 : Metabolic pathways | 153 | 1334 | 0.00001 |
| F2 | T vs stage IV | P4-phase | Mineral Absorption | ko04978 : Mineral absorption | 6 | 108 | 0.0026 |
| F2 | T vs stage IV | P4-phase | Cell death | ko04217 : Necroptosis | 7 | 228 | 0.0137 |
| F2 | T vs stage IV | P4-phase | Cell death | ko04216 : Ferroptosis | 5 | 107 | 0.0137 |
| F2 | T vs stage IV | P4-phase | Other metabolisms | ko04141 : Protein processing in endoplasmic reticulum | 8 | 205 | 0.0026 |
| F2 | T vs stage I | T-phase | **Cancer related pathways** | ko05217 : Basal cell carcinoma | 3 | 70 | 0.0265 |
| F2 | T vs stage I | T-phase | **Cancer related pathways** | ko05200 : Pathways in cancer | 6 | 584 | 0.0490 |
| F2 | T vs stage I | T-phase | Steroid metabolism | ko04934 : Cushing syndrome | 4 | 162 | 0.0265 |
| F2 | T vs stage I | T-phase | Endocrine system (non-steroidal) | ko04916 : Melanogenesis | 3 | 110 | 0.0490 |
